# A robust phylogenomic timetree for biotechnologically and medically important fungi in the genera *Aspergillus* and *Penicillium*

**DOI:** 10.1101/370429

**Authors:** Jacob L. Steenwyk, Xing-Xing Shen, Abigail L. Lind, Gustavo H. Goldman, Antonis Rokas

## Abstract

**Abbreviations:** NT, nucleotide; AA, amino acid; CI, credible interval; RCV, relative composition variability; IC, internode certainty; GSF, gene support frequencies; GLS, gene-wise log-likelihood scores; DVMC, degree of violation of a molecular clock

The filamentous fungal family Aspergillaceae contains > 1,000 known species, mostly in the genera *Aspergillus* and *Penicillium*. Several species are used in the food, biotechnology, and drug industries (e.g., *Aspergillus oryzae, Penicillium camemberti*), while others are dangerous human and plant pathogens (e.g., *Aspergillus fumigatus, Penicillium digitatum*). To infer a robust phylogeny and pinpoint poorly resolved branches and their likely underlying contributors, we used 81 genomes spanning the diversity of *Aspergillus* and *Penicillium* to construct a 1,668-gene data matrix. Phylogenies of the nucleotide and amino acid versions of this full data matrix as well as of five additional 834-gene data matrices constructed by subsampling the top 50% of genes according to different criteria associated with strong phylogenetic signal were generated using three different maximum likelihood schemes (i.e., gene-partitioned, unpartitioned, and coalescence). Examination of the topological agreement among these 36 phylogenies and measures of internode certainty identified 12 / 78 (15.4%) bipartitions that were incongruent and pinpoint the likely underlying contributing factors (incomplete lineage sorting, hybridization or introgression, and reconstruction artifacts associated with poor taxon sampling). Relaxed molecular clock analyses suggest that Aspergillaceae likely originated in the lower Cretaceous and the *Aspergillus* and *Penicillium* genera in the upper Cretaceous. Our results shed light on the ongoing debate on *Aspergillus* systematics and taxonomy and provide a robust evolutionary and temporal framework for comparative genomic analyses in Aspergillaceae. More broadly, our approach provides a general template for phylogenomic identification of resolved and contentious branches in densely genome-sequenced lineages across the tree of life.

## Introduction

The vast majority of the 1,062 described species from the family Aspergillaceae (phylum Ascomycota, class Eurotiomycetes, order Eurotiales) (Houbraken et al. 2014) belong to the genera *Aspergillus* (42.5%; 451 / 1,062) and *Penicillium* (51.6%; 549 / 1,062) (Benson et al. 2007; Sayers et al. 2009). Fungi from Aspergillaceae exhibit diverse ecologies; for example, *Penicillium verrucosum* is widespread in cold climates but has yet to be isolated in the tropics (Pitt 2002), whereas *Aspergillus nidulans* is able to grow at a wide range of temperatures but favors warmer ones (Ogundero 1983). Several representative species in the family are exploited by humans, while a number of others are harmful to humans or their activities (Gibbons and Rokas 2013). Examples of useful-to-humans organisms among *Aspergillus* species include *Aspergillus oryzae*, which is used in the production of traditional Japanese foods including soy sauce, sake, and vinegar (Machida et al. 2008; Gibbons et al. 2012) as well as of amylases and proteases (Kobayashi et al. 2007) and *Aspergillus terreus*, which produces mevinolin (lovastatin), the potent cholesterol-lowering drug (Albert et al. 1980). Examples of useful-to-humans *Penicillium* species include *Penicillium camemberti* and *Penicillium roqueforti*, which contribute to cheese production (Nelson 1970; Lessard et al. 2012), and *Penicillium citrinum*, which produces the cholesterol lowering drug mevastatin, the world’s first statin (Endo 2010). In contrast, examples of harmful-to-humans organisms include the pathogen, allergen, and mycotoxin-producing species *Aspergillus fumigatus* and *Aspergillus flavus* (Nierman et al. 2005; Hedayati et al. 2007) and the post-harvest pathogens of citrus fruits, stored grains, and other cereal crops *Penicillium expansum, Penicillium digitatum,* and *Penicillium italicum* (Marcet-Houben et al. 2012; Ballester et al. 2015; Li et al. 2015).

Much of the ubiquity, ecological diversity, and wide impact on human affairs that Aspergillaceae exhibit is reflected in their phenotypic diversity, including their extremotolerance (e.g., ability to withstand osmotic stress and wide temperature range) (Magan and Lacey 1984; Marín et al. 1998; Pitt and Hocking 2009; Vinnere Pettersson and Leong 2011) and ability to grown on various carbon sources (Pitt and Hocking 2009; de Vries et al. 2017). Fungi from Aspergillaceae are also well known for their ability to produce a remarkable diversity of secondary metabolites, small molecules that function as toxins, signaling molecules, and pigments (Pitt 1994; Keller et al. 2005; Frisvad and Larsen 2015; Macheleidt et al. 2016; Rokas et al. 2018). Secondary metabolites likely play key roles in fungal ecology (Rohlfs et al. 2007; Fox and Howlett 2008; Stierle and Stierle 2015), but these small molecules often have biological activities that are either harmful or beneficial to human welfare. For example, the *A. fumigatus*-produced secondary metabolite gliotoxin is a potent virulence factor in cases of systemic mycosis in vertebrates (Rohlfs and Churchill 2011), and the *A. flavus*-produced secondary metabolite aflatoxin is among the most toxic and carcinogenic naturally occurring compounds (Squire 1981; Keller et al. 2005). In contrast, other secondary metabolites are mainstay antibiotics and pharmaceuticals; for example, the *Penicillium chrysogenum*produced penicillin is among the world’s most widely used antibiotics (Chain et al. 1940; Fleming 1980; Aminov 2010) and the *P. citrinum*-produced cholesterol lowering statins are consistently among the world’s blockbuster drugs (Endo 2010).

Understanding the evolution of the diverse ecological lifestyles exhibited by Aspergillaceae members as well as the family’s remarkable chemodiversity requires a robust phylogenetic framework. To date, most molecular phylogenies of the family Aspergillaceae are derived from single or few genes and have yielded conflicting results. For example, it is debated whether the genus *Aspergillus* is monophyletic or if it includes species from other genera such as *Penicillium* (Pitt and Taylor 2014; Samson et al. 2014). Furthermore, studies using genome-scale amounts of data, which could have the power to resolve evolutionary relationships and identify underlying causes of conflict (Rokas et al. 2003; Salichos and Rokas 2013), have so far tended to use a small subset of fungi from either *Aspergillus* or *Penicillium* (Nielsen et al. 2017; de Vries et al. 2017; Kjærbølling et al. 2018). Additionally, these genome-scale studies do not typically examine the robustness of the produced phylogeny; rather, based on the high clade support values (e.g., bootstrap values) obtained, these studies infer that the topology obtained is highly accurate (Yang et al. 2016; Nielsen et al. 2017; de Vries et al. 2017; Kjærbølling et al. 2018).

In recent years, several phylogenomic analyses have shown that high clade support values can be misleading (Phillips et al. 2004; Kumar et al. 2012; Salichos and Rokas 2013), that incongruence, the presence of topological conflict between different data sets or analyses, is widespread (Hess and Goldman 2011; Song et al. 2012; Salichos and Rokas 2013; Zhong et al. 2013), and that certain branches of the tree of life can be very challenging to resolve, even with genome-scale amounts of data (Shen et al. 2016; Suh 2016; Arcila et al. 2017; King and Rokas 2017; Shen et al. 2017). Comparison of the topologies inferred in previous phylogenomic studies in Aspergillaceae (Yang et al. 2016; Nielsen et al. 2017; de Vries et al. 2017; Kjærbølling et al. 2018) suggests the presence of incongruence (Figure S1). For example, some studies have reported section *Nidulantes* to be the sister group to section *Nigri* (de Vries et al. 2017), whereas other studies have placed it as the sister group to *Ochraceorosei* (Kjærbølling et al. 2018) (Figure S1).

A robust phylogeny of Aspergillaceae is also key to establishing a robust taxonomic nomenclature for the family. In recent years, the taxonomy of *Aspergillus* and *Penicillium* has been a point of contention due to two key differences among inferred topologies based on analyses of a few genes (Kocsubé et al. 2016; Taylor et al. 2016). The first key difference concerns the placement of the genus *Penicillium*. One set of analyses places the genus as a sister group to *Aspergillus* section *Nidulantes*, which would imply that *Penicillium* is a section within the genus *Aspergillus* (Taylor et al. 2016), whereas a different set of analyses suggests that the genera *Penicillium* and *Aspergillus* are reciprocally monophyletic (Kocsubé et al. 2016). The second key difference concerns whether sections *Nigri*, *Ochraceorosei*, *Flavi*, *Circumdati*, *Candidi*, and *Terrei*, which are collectively referred to as “narrow *Aspergillus*”, form a monophyletic group (Taylor et al. 2016) or not (Kocsubé et al. 2016). Both of these differences are based on analyses of a few genes (4 loci, Taylor et al. 2016 and 9 loci, Kocsubé et al. 2016) and the resulting phylogenies typically exhibit low support values for deep internodes, including for the ones relevant to this debate.

To shed light on relationships among these fungi, we employed a genome-scale approach to infer the evolutionary history among *Aspergillus*, *Penicillium*, and other fungal genera from the family Aspergillaceae. More specifically, we used the genome sequences of 81 fungi from Aspergillaceae spanning 5 genera, 25 sections within *Aspergillus* and *Penicillium*, and 12 outgroup fungi to construct nucleotide (NT) and amino acid (AA) versions of a data matrix comprised of 1,668 orthologous genes. Using three different maximum likelihood schemes (i.e., gene-partitioned, unpartitioned, and coalescence), we inferred phylogenies from the 1,668-gene data matrix as well as from five additional 834-gene data matrices derived from the top 50% of genes harboring strong phylogenetic signal according to five different criteria (alignment length, average bootstrap value, taxon completeness, treeness / relative composition variability, and number of variable sites). Comparisons of these phylogenies coupled with complementary measures of internode certainty (Salichos and Rokas 2013; Salichos et al. 2014; Kobert et al. 2016) identified 12 / 78 (15.4%) incongruent bipartitions in the phylogeny of Aspergillaceae. These cases of incongruence can be grouped into three categories: (i) 2 shallow bipartitions with low levels of incongruence likely driven by incomplete lineage sorting, (ii) 3 shallow bipartitions with high levels of incongruence likely driven by hybridization or introgression (or very high levels of incomplete lineage sorting), and (iii) 7 deeper bipartitions with varying levels of incongruence likely driven by reconstruction artifacts likely linked with poor taxon sampling. We also estimated divergence times across Aspergillaceae using relaxed molecular clock analyses. Our results suggest Aspergillaceae originated in the lower Cretaceous, 117.4 (95% Credible Interval (CI): 141.5 - 96.9) million years ago (mya), and that *Aspergillus* and *Penicillium* originated 81.7 mya (95% CI: 87.5 - 72.9) and 73.6 mya (95% CI: 84.8 - 60.7), respectively. We believe this phylogeny and timetree are highly informative with respect to the ongoing debate on *Aspergillus* systematics and taxonomy, and provide a state-of-the-art platform for comparative genomic, ecological, and chemodiversity studies in this ecologically diverse and biotechnologically and medically significant family of filamentous fungi.

## Methods

### Genome sequencing and assembly

Mycelia were grown on potato dextrose agar for 72 hours before lyophilization. Lyophilized mycelia were lysed by grinding in liquid nitrogen and suspension in extraction buffer (100 mM Tris-HCl pH 8, 250 mM NaCl, 50 mM EDTA, and 1% SDS). Genomic DNA was isolated from the lysate with a phenol/chloroform extraction followed by an ethanol precipitation.

DNA was sequenced with both paired-end and mate-pair strategies to generate a high-quality genome assembly. Paired-end libraries and Mate-pair libraries were constructed at the Genomics Services Lab at HudsonAlpha (Huntsville, Alabama) and sequenced on an Illumina HiSeq X sequencer. Paired-end libraries were constructed with the Illumina TruSeq DNA kit, and mate-pair libraries were constructed with the Illumina Nextera Mate Pair Library kit targeting an insert size of 4 Kb. In total, 63 million paired-end reads and 105 million mate-pair reads, each of which was 150 bp in length, were generated.

The *A. spinulosporus* genome was assembled using the iWGS pipeline (Zhou et al. 2016). Paired-end and mate-pair reads were assembled with SPADES, version 3.6.2 (Bankevich et al. 2012), using optimal k-mer lengths chosen using KMERGENIE, version 1.6982 (Chikhi and Medvedev 2014) and evaluated with QUAST, version 3.2 (Gurevich et al. 2013). The resulting assembly is 33.8 MB in size with an N50 of 939 Kb.

### Data collection and quality assessment

To collect a comprehensive set of genomes representative of Aspergillaceae, we used “Aspergillaceae” as a search term in NCBI’s Taxonomy Browser and downloaded a representative genome from every species that had a sequenced genome as of February 5^th^ 2018. We next confirmed that each species belonged to Aspergillaceae according to previous literature reports (Houbraken and Samson 2011; de Vries et al. 2017). Altogether, 80 publicly available genomes and 1 newly sequenced genome spanning 5 genera (45 *Aspergillus* species; 33 *Penicillium* species; one *Xeromyces* species; one *Monascus* species; and one *Penicilliopsis* species) from the family Aspergillaceae were collected (File S1). We also retrieved an additional 12 fungal genomes from representative species in the order Eurotiales but outside the family Aspergillaceae to use as outgroups.

To determine if the genomes contained gene sets of sufficient quality for use in phylogenomic analyses, we examined their gene set completeness using Benchmarking Universal Single-Copy Orthologs (BUSCO), version 2.0.1 (Waterhouse et al. 2018) (Figure S2). In brief, BUSCO uses a consensus sequence built from hidden Markov models derived from 50 different fungal species using HMMER, version 3.1b2 (Eddy 2011) as a query in TBLASTN (Camacho et al. 2009; Madden 2013) to search an individual genome for 3,156 predefined orthologs (referred to as BUSCO genes) from the Pezizomycotina database (creation date: 02-13-2016) available from ORTHODB, version 9 (Waterhouse et al. 2013). To determine the copy number and completeness of each BUSCO gene in a genome, gene structure is predicted using AUGUSTUS, version 2.5.5 (Stanke and Waack 2003), from the nucleotide coordinates of putative genes identified using BLAST and then aligned to the HMM alignment of the same BUSCO gene. Genes are considered “single copy” if there is only one complete predicted gene present in the genome, “duplicated” if there are two or more complete predicted genes for one BUSCO gene, “fragmented” if the predicted gene is shorter than 95% of the aligned sequence lengths from the 50 different fungal species, and “missing” if there is no predicted gene.

### Phylogenomic data matrix construction

In addition to their utility as a measure of genome completeness, BUSCO genes have also proven to be useful markers for phylogenomic inference (Waterhouse et al. 2018), and have been successfully used in phylogenomic studies of clades spanning the tree of life, such as insects (Ioannidis et al. 2017) and budding yeasts (Shen et al. 2016). To infer evolutionary relationships, we constructed nucleotide (NT) and amino acid (AA) versions of a data matrix comprised of the aligned and trimmed sequences of numerous BUSCO genes (Figure S3). To construct this data matrix, we first used the BUSCO output summary files to identify orthologous single copy BUSCO genes with > 50% taxon-occupancy (i.e., greater than 47 / 93 taxa have the BUSCO gene present in their genome); 3,138 (99.4%) BUSCO genes met this criterion. For each BUSCO gene, we next created individual AA fasta files by combining sequences across all taxa that have the BUSCO gene present. For each gene individually, we aligned the sequences in the AA fasta file using MAFFT, version 7.294b (Katoh and Standley 2013), with the BLOSUM62 matrix of substitutions (Mount 2008), a gap penalty of 1.0, 1,000 maximum iterations, and the “genafpair” parameter. To create a codon-based alignment, we used a custom PYTHON, version 3.5.2 (https://www.python.org/), script using BIOPYTHON, version 1.7 (Cock et al. 2009), to thread codons onto the AA alignment. The NT and AA sequences were then individually trimmed using TRIMAL, version 1.4 (Capella-Gutierrez et al. 2009), with the “automated1” parameter. We next removed BUSCO genes whose sequence lengths were less than 50% of the untrimmed length in either the NT or AA sequences resulting in 1,773 (56.2%) BUSCO genes. Lastly, we removed BUSCO genes whose trimmed sequence lengths were too short (defined as genes whose alignment length was less than or equal to 167 AAs and 501 NTs), resulting in 1,668 (52.9%) BUSCO genes. The NT and AA alignments of these 1,668 BUSCO genes were then concatenated into the full 1,668-gene NT and AA versions of the phylogenomic data matrix.

To examine the stability of inferred relationships across all taxa, we constructed additional NT and AA data matrices by subsampling genes from the 1,668-gene data matrix that harbor signatures of strong phylogenetic signal. More specifically, we used 5 measures associated with strong phylogenetic signal (Shen et al. 2016) to create 5 additional data matrices (1 data matrix per measure) comprised of the top scoring 834 (50%) genes for NTs and AAs (Figure S4). These five measures were: alignment length, average bootstrap value, taxon completeness, treeness / relative composition variability (RCV) (Phillips and Penny 2003), and the number of variable sites. We calculated each measure with custom PYTHON scripts using BIOPYTHON. Treeness / RCV was calculated using the following formula:

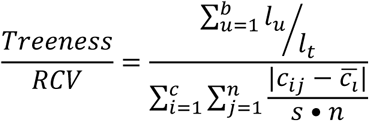

where *l*_*u*_ refers to the internal branch length of the *u*th branch (of *b* internal branches), *l*_*t*_ refers to total tree length, *c* is the number of different characters per sequence type (4 for nucleotides and 20 for amino acids), *n* is the number of taxa in the alignment, *c*_*ij*_ refers to the number of *i*th *c* characters for the *j*th taxon, 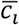 refers to the average number of the *i*th *c* character across *n* taxa, and *s* refers to the total number of sites in the alignment. Altogether, we constructed a total of 12 data matrices (one 1,668-gene NT data matrix, one 1,668-gene AA data matrix, five NT subsample data matrices, and five AA subsample data matrices).

### Maximum likelihood phylogenetic analyses

We implemented a maximum likelihood framework to infer evolutionary relationships among taxa for each of the 1,668 single genes and each of the 12 data matrices separately. For inferences made using either the 1,668- or 834-gene data matrices, we used three different analytical schemes: concatenation with gene-based partitioning, concatenation without partitioning, and gene-based coalescence (Felsenstein 1981; Rokas et al. 2003; Edwards 2009; Mirarab and Warnow 2015). All phylogenetic trees were built using IQ-TREE, version 1.6.1 (Nguyen et al. 2015). In each case, we determined the best model for each single gene or partition using the “-m TEST” and “-mset raxml” parameters, which automatically estimate the best fitting model of substitutions according to their Bayesian Information Criterion values for either NTs or AAs (Kalyaanamoorthy et al. 2017) for those models shared by RAXML (Stamatakis 2014) and IQ-TREE.

We first examined the inferred best fitting models across all single gene trees. Among NT genes, the best fitting model for 1,643 genes was a general time reversible model with unequal rates and unequal base frequencies with discrete gamma models, “GTR+G4” (Tavaré 1986; Yang 1994; Yang 1996), and for the remaining 25 genes was a general time reversible model with invariable sites plus discrete gamma models, “GTR+I+G4” (Tavaré 1986; Vinet and Zhedanov 2011) (Figure S5a). Among AA genes, the best fitting model for 643 genes was the JTT model with invariable sites plus discrete gamma models, “JTT+I+G4” (Jones et al. 1992; Vinet and Zhedanov 2011), for 362 genes was the LG model with invariable sites and discrete gamma models, “LG+I+G4” (Le and Gascuel 2008; Vinet and Zhedanov 2011), for 225 genes was the JTT model with invariable sites, empirical AA frequencies, and discrete gamma models “JTT+F+I+G4” (Jones et al. 1992; Vinet and Zhedanov 2011), and for 153 genes was the JTTDCMut model with invariable sites and discrete gamma models, “JTTDCMut+I+G4” (Kosiol and Goldman 2005; Vinet and Zhedanov 2011) (Figure S5b). We used IQ-TREE for downstream analysis because a recent study using diverse empirical phylogenomic data matrices showed that it is a top-performing software (Zhou et al. 2018).

To reconstruct the phylogeny of Aspergillaceae using a partitioned scheme where each gene has its own model of sequence substitution and rate heterogeneity across sites parameters for any given data matrix, we created an additional input file describing these and gene boundary parameters. More specifically, we created a nexus-format partition file that was used as input with the “-spp” parameter, which allows each gene partition in the data matrix to have its set of evolutionary rates (Chernomor et al. 2016). To increase the number of candidate trees used during maximum likelihood search, we changed the “-nbest” parameter from the default value of 5 to 10. Lastly, we conducted 5 independent searches for the maximum likelihood topology using 5 distinct seeds specified with the “-seed” parameter and chose the search with the best log-likelihood score. We used the phylogeny inferred using a partitioned scheme on the full NT data matrix as the reference one for all subsequent comparisons (Figure 1).

**Figure 1.**
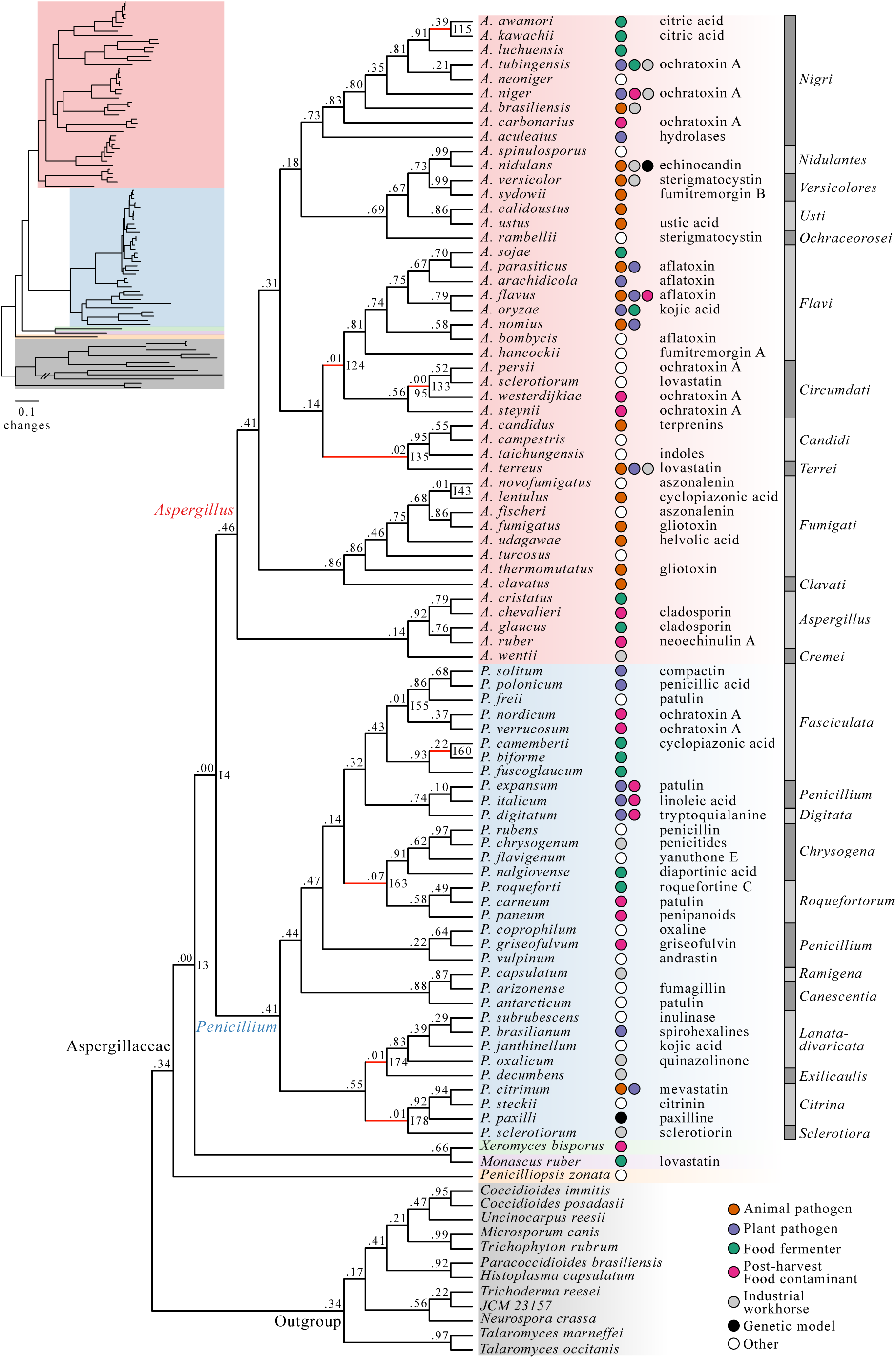
A robust genome-scale phylogeny for the fungal family Aspergillaceae. Different genera are depicted using different colored boxes; *Aspergillus* is shown in red, *Penicillium* in blue, *Xeromyces* in green, *Monascus* in purple, and *Penicilliopsis* in orange. Different sections within *Aspergillus* and *Penicillium* are depicted with alternating dark grey and grey bars. Internode certainty values are shown below each internode and bootstrap values are shown above each internode (only bootstrap values lower than 100 percent are shown). Internode certainty values were calculated using the 1,668 maximum likelihood single gene trees. 5,000 ultrafast bootstrap replicates were used to determine internode support. Internodes were considered unresolved if they were not present in one or more of the other 35 phylogenies represented in Figure 2 – the branches of these unresolved internodes are drawn in red. The inset depicts the phylogeny with branch lengths corresponding to estimated nucleotide substitutions per site. Colored circles next to species names indicate the lifestyle or utility of the species (i.e., animal pathogen, dark orange; plant pathogen, purple; food fermenter, green; post-harvest food contaminant, pink; industrial workhorse, grey; genetic model, black; other, white). Exemplary secondary metabolites produced by different Aspergillaceae species are written to the right of the colored circles.

To infer the phylogeny of Aspergillaceae using a non-partitioned scheme, we used a single model of sequence substitution and rate heterogeneity across sites for the entire matrix. To save computation time, the most appropriate single model was determined by counting which best fitting model was most commonly observed across single gene trees. The most commonly observed model was “GTR+F+I+G4” (Waddell and Steel 1997; Vinet and Zhedanov 2011), which was favored in 1,643 / 1,668 (98.5%) of single genes, and “JTT+I+G4” (Jones et al. 1992; Vinet and Zhedanov 2011), which was favored in 643 / 1,668 (38.5%) of single genes, for NTs and AAs, respectively, (Figure S5). In each analysis, the chosen model was specified using the “-m” parameter.

To reconstruct the phylogeny of Aspergillaceae using coalescence, a method that estimates species phylogeny from single gene trees under the multi-species coalescent (Edwards 2009), we combined all NEWICK (Felsenstein 1986; Felsenstein 1996) formatted single gene trees inferred using their best fitting models into a single file. The resulting file was used as input to ASTRAL-II, version 4.10.12 (Mirarab and Warnow 2015) with default parameters.

To evaluate support for single gene trees and for the reference phylogeny (Figure 1), we used the ultrafast bootstrap approximation approach (UFBoot) (Hoang et al. 2018), an accurate and faster alternative to the classic bootstrap approach. To implement UFBoot for the NT 1,668-gene data matrix and single gene trees, we used the “-bb” option in IQ-TREE with 5,000 and 2,000 ultrafast bootstrap replicates, respectively.

### Evaluating topological support

To identify and quantify incongruence, we used two approaches. In the first approach, we compared the 36 topologies inferred from the full 1,668-gene NT and AA data matrices and five additional 834-gene data matrices (constructed by selecting the genes that have the highest scores in five measures previously shown to be associated with strong phylogenetic signal; see above) using three different maximum likelihood schemes (i.e., gene partitioned, non-partitioned, coalescence) and identified all incongruent bipartitions between the reference phylogeny (Figure 1) and the other 35. In the second approach, we scrutinized each bipartition in the reference phylogeny using measures of internode certainty (IC) measures for complete and partial single gene trees (Salichos and Rokas 2013; Salichos et al. 2014; Kobert et al. 2016). To better understand single gene support among conflicting bipartitions, we calculated gene-wise log-likelihood scores (GLS) (Shen et al. 2017) and gene support frequencies (GSF) for the reference and alternative topologies at conflicting bipartitions.

#### Identifying internodes with conflict across subsampled data matrices

To identify incongruent bipartitions between the reference phylogeny and the other 35 phylogenies, we first included the 36 generated phylogenetic trees into a single file. We next evaluated the support of all bipartitions in the reference topology among the other 35 phylogenies using the “-z” option in RAXML. Any bipartition in the reference phylogeny that was not present in the rest was considered incongruent; each conflicting bipartition was identified through manual examination of the conflicting phylogenies. To determine if sequence type, subsampling method, or maximum likelihood scheme was contributing to differences in observed topologies among conflicting internodes, we conducted multiple correspondence analysis of these features among the 36 phylogenies and visualized results using the R, version 3.3.2 (R Development Core Team 2008), packages FACTOMINER, version 1.40 (Lê et al. 2008) and FACTOEXTRA, version 1.0.5 (Kassambara and Mundt 2017).

#### Identifying internodes with conflict across the 1,668 gene trees

To examine the presence and degree of support of conflicting bipartitions, we calculated the internode certainty (Salichos and Rokas 2013; Salichos et al. 2014; Kobert et al. 2016; Zhou et al. 2017) of all internodes in the reference phylogeny (Figure 1) using the 1,668 gene trees as input. In general, IC scores near 0 indicate that there is near-equal support for an alternative, conflicting bipartition among a set of trees compared to a given bipartition present in the reference topology, which is indicative of high conflict. Therefore, we investigated incongruence in all internodes in the reference phylogeny (Figure 1) that exhibited IC scores lower than 0.1. To calculate IC values for each bipartition for the reference phylogeny, we created a file with all 1,668 complete and partial single gene trees. The resulting file of gene trees, specified with the “-z” parameter in RAXML, were used to calculate IC values using the “-f i” argument. The topology was specified with the “-t” parameter. Lastly, we used the Lossless corrected IC scoring scheme, which corrects for variation in taxon number across single gene trees (Kobert et al. 2016). We also used these IC values to inform which data type (NT or AA) provided the strongest signal for the given set of taxa and sequences. We observed that NTs consistently exhibited higher IC scores than AAs (hence our decision to use the topology inferred from the full NT data matrix using a gene-partitioned scheme – shown in Figure 1 – as the “reference” topology in all downstream analyses).

#### Examining gene-wise log-likelihood scores for incongruent internodes

To determine the per gene distribution of phylogenetic signal supporting a bipartition in the reference phylogeny or a conflicting bipartition, we calculated gene-wise log-likelihood scores (GLS) (Shen et al. 2017) using the NT data matrix. We chose to calculate GLS using the NT data matrix because distributions of IC values from phylogenies inferred using NTs had consistently higher IC values across schemes and data matrices (Figure S6). To do so, we used functions available in IQ-TREE. More specifically, we inputted a phylogeny with the reference or alternative topology using the “-te” parameter and informed IQ-TREE of gene boundaries, their corresponding models, and optimal rate heterogeneity parameters in the full 1,668-gene data matrix using the “-spp” parameter. Lastly, we specified that partition log-likelihoods be outputted using the “-wpl” parameter. To determine if a gene provided greater support for the reference or alternative bipartition, we calculated the difference in GLS (∆GLS) using the following formula:

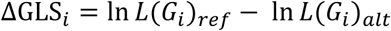

where *ln L*(*G*_*i*_)_*ref*_ and *ln L*(*G*_*i*_)_*alt*_ represent the log-likelihood values for the reference and alternative topologies for gene *G*_*i*_. Thus, values greater than 0 reflect genes in favor of the reference bipartition, values lower than 0 reflect genes in favor of the alternative bipartition, and values of 0 reflect equal support between the reference and alternative bipartitions.

#### Calculating gene support frequencies for reference and conflicting bipartitions

We next examined support for bipartitions in the reference topology as well as for their most prevalent conflicting bipartitions by calculating their gene support frequencies (GSF). GSF refers to the fraction of single gene trees that recover a particular bipartition. Currently, RAXML can only calculate GSF for trees with full taxon representation. Since our dataset contained partial gene trees, we conducted customs tests for determining GSF. To calculate GSF for NT (GSF_NT_) and AA (GSF_AA_) single gene trees, we extracted subtrees for the taxa of interest in individual single gene trees and counted the occurrence of various topologies. For example, consider there are three taxa represented as A, B, and C, the reference rooted topology is “((A,B),C);” and the alternative rooted topology is “((A,C),B);”. We counted how many single gene trees supported “(A,B),” or “(A, C),”. For reference and alternative topologies involving more than three taxa or sections, we conducted similar tests. For example, if the reference rooted topology is “(((A,B),C),D);” and the alternative rooted topology is “((A,B),(C,D));”, we counted how many single gene phylogenies supported “((A,B),C),” as sister to D and how many single gene phylogenies supported “(A,B),” and “(C,D),” as pairs of sister clades. For conflicting bipartitions at shallow depths in the phylogeny (i.e., among closely related species), we required all taxa to be present in a single gene tree; for conflicting bipartitions near the base of the phylogeny (i.e., typically involving multiple sections), we required at least one species to be present from each section of interest. Scripts to determine GSF were written using functions provided in NEWICK UTILITIES, version 1.6 (Junier and Zdobnov 2010).

#### Topology tests

To test the previously reported hypotheses of a) the genus *Penicillium* being the sister group to *Aspergillus* section *Nidulantes* and b) monophyly of narrow *Aspergillus* (sections *Nigri*, *Ochraceorosei*, *Flavi*, *Circumdati*, *Candidi*, *Terrei*) (Taylor et al. 2016), we conducted a series of tree topology tests using the 1,668-gene nucleotide data matrix using IQ-TREE (Nguyen et al. 2015). More specifically, we used the “GTR+F+I+G4” model and conducted the Shimodaira-Hasegawa (Shimodaira and Hasegawa 1999) and the approximately unbiased tests (Shimodaira 2002) as specified with the “-au” parameter. These tests were conducted using 10,000 resamplings using the resampling estimated log-likelihood (RELL) method (Kishino et al. 1990) as specified by the “-zb” parameter. We tested each hypothesis separately by generating the maximum likelihood topology under the constraint that the hypothesis is correct (specified using the “-z” parameter) and comparing its likelihood score to the score of the unconstrained maximum likelihood topology.

### Estimating divergence times

To estimate the divergence times for the phylogeny of the Aspergillaceae, we analyzed our NT data matrix used the Bayesian method implemented in MCMCTREE from the PAML package, version 4.9d (Yang 2007). To do so, we conducted four analyses: we (i) identified genes evolving in a “clock-like” manner from the full data matrix, (ii) estimated the substitution rate across these genes, (iii) estimated the gradient and Hessian (Dos Reis and Yang 2013) at the maximum likelihood estimates of branch lengths, and (iv) estimated divergence times by Markov chain Monte Carlo (MCMC) analysis.

#### (i) Identifying “clock-like” genes

Currently, large phylogenomic data matrices that contain hundreds to thousands of genes and many dozens of taxa are intractable for Bayesian inference of divergence times; thus, we identified and used only those genes that appear to have evolved in a “clock-like” manner in the inference of divergence times. To identify genes evolving in a “clock-like” manner, we calculated the degree of violation of a molecular clock (DVMC) (Liu et al. 2017) for single gene trees. DVMC is the standard deviation of root to tip distances in a phylogeny and is calculated using the following formula:

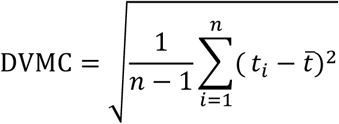

where *t*_*i*_ represents the distance between the root and species *i* across *n* species. Using this method, genes with low DVMC values evolve in a “clock-like” manner compared to those with higher values. We took the top scoring 834 (50%) genes to estimate divergence times.

#### (ii) Estimating substitution rate

To estimate the substitution rate across the 834 genes, we used BASEML from the PAML package, version 4.9d (Yang 2007). We estimated substitution rate using a “GTR+G” model of substitutions (model = 7) and a strict clock model (clock = 1). Additionally, we point calibrated the root of the tree to 96 million years ago (mya) according to TIMETREE (Hedges et al. 2006), which is based on several previous estimates (Berbee and Taylor 2001: 50.0 mya; Vijaykrishna et al. 2006: 96.1 mya; Sharpton et al. 2009: 146.1 mya). We estimated a substitution rate of 0.04 substitutions per 10 million years.

#### (iii) Estimation of the gradient and Hessian

To save computing time, the likelihood of the alignment was approximated using a gradient and Hessian matrix. The gradient and Hessian refer to the first and second derivatives of the log-likelihood function at the maximum likelihood estimates of branch lengths (Dos Reis and Yang 2013), and collectively describe the curvature of the log-likelihood surface. Estimating gradient and Hessian requires an input tree with specified time constraints. For time constraints, we used the *Aspergillus flavus – Aspergillus oryzae* split (3.68-3.99 mya: Sharpton et al. 2009; Da Lage et al. 2013), the *Aspergillus fumigatus – Aspergillus clavatus* split (35-59 mya: Sharpton et al. 2009; Da Lage et al. 2013), the origin of the genus *Aspergillus* (43-85 mya: Kensche et al. 2008; Sharpton et al. 2009; Beimforde et al. 2014; Fan et al. 2015; Gaya et al. 2015), and the origin of Aspergillaceae (50-146 mya: Berbee and Taylor 2001; Vijaykrishna et al. 2006; Sharpton et al. 2009) as obtained from TIMETREE (Hedges et al. 2006).

#### (iv) Estimating divergence times using MCMC analysis

To estimate divergence times using a relaxed molecular clock (clock = 2), we used the resulting gradient and Hessian results from the previous step for use in MCMC analysis using MCMCTREE (Yang 2007) and the topology inferred using the gene partitioned approach and the 834-gene NT matrix from the top scoring DVMC genes. To do so, a gamma distribution prior shape and scale must be specified. The gamma distribution shape and scale is determined from the substitution rate determined in step ii where shape is *a*=(*s*/*s*)^*2*^ and scale is *b*=*s*/*s*^*2*^ and s is the substitution rate. Therefore, *a*=*1* and *b*=*25* and the “rgene_gamma” parameter was set to “1 25.” We also set the “sigma2_gamma” parameter to “1 4.5.” To minimize the effect of initial values on the posterior inference, we discarded the first 100,000 results. Thereafter, we sampled every 500 iterations until 10,000 samples were gathered. Altogether, we ran 5.1 million iterations (100,000 + 500 × 10,000), which is 510 times greater than the recommended minimum for MCMC analysis (Raftery and Lewis 1995). Lastly, we set the “finetune” parameter to 1.

### Statistical analysis and figure making

All statistical analyses were conducted in R, version 3.3.2 (R Development Core Team 2008). Spearman rank correlation analyses (Sedgwick 2014) were conducted using the “rcorr” function in the package HMISC, version 4.1-1 (Harrell Jr 2015). Stacked barplots, barplots, histograms, scatterplots, and boxplots were made using GGPLOT2, version 2.2.1 (Wickham 2009). Intersection plots (also known as UpSet plots), were made using UPSETR, version 1.3.3 (Conway et al. 2017). The topological similarity heatmap and hierarchical clustering were done using PHEATMAP, version 1.0.8 (Kolde 2012). Phylogenetic trees were visualized using FIGTREE, version 1.4.3 (Rambaut 2009). The phylogenetic tree with the geological time scale was visualized using STRAP, version 1.4 (Bell and Lloyd 2015). Artistic features of figures (e.g., font size, font style, etc.) were minimally edited using the graphic design software Affinity Designer (https://affinity.serif.com/enus/).

## Results

### The examined genomes have nearly complete gene sets

Assessment of individual gene set completeness showed that most of the 93 genomes (81 in the ingroup and 12 in the outgroup) used in our study contain nearly complete gene sets and that all 93 genomes are appropriate for phylogenomic analyses. Specifically, the average percentage of BUSCO single-copy genes from the Pezizomycotina database (Waterhouse et al. 2013) present was 96.2 ± 2.6% (minimum: 81.1%; maximum: 98.9%; Figure S2). Across the 93 genomes, only 3 (3.2%) genomes had < 90% of the BUSCO genes present in single-copy (*Penicillium carneum*: 88.6%; *Penicillium verrucosum*: 86.1%; and *Histoplasma capsulatum*: 81.1%).

### The generated data matrices exhibit very high taxon occupancy

The NT and AA alignments of the 1,668-gene data matrix were comprised of 3,163,258 and 1,054,025 sites, respectively. The data matrix exhibited very high taxon occupancy (average gene taxon occupancy: 97.2 ± 0.1%; minimum: 52.7%; maximum: 100%; Figure S7a, b; File S2). 417 genes had 100% taxon-occupancy, 1,176 genes had taxon-occupancy in the 90% to 99.9% range, and only 75 genes had taxon occupancy lower than 90%. Assessment of the 1,668 genes for five criteria associated with strong phylogenetic signal (gene-wise alignment length, average bootstrap value, completeness, treeness / RCV, and the number of variable sites) facilitated the construction of five subsampled matrices derived from 50% of the top scoring genes (Figure S7; File S2).

Examination of the gene content differences between the 5 NT subsampled data matrices as well as between the 5 AA data matrices revealed that they are comprised of variable sets of genes (Figure S8). For example, the largest intersection among NT data matrices comprised of 207 genes that were shared between all NT matrices except the completeness-based one; similarly, the largest intersection among AA data matrices was 228 genes and was shared between all AA matrices except the completeness-based one (Figure S8a, b). Examination of the number of gene overlap between the NT and AA data matrices for each criterion (Figure S8c) showed that three criteria yielded identical or nearly identical NT and AA gene sets. These were completeness (834 / 834; 100% shared genes; *r*_*s*_ = 1.00, *p* < 0.01; Figure S7c), alignment length (829 / 834; 99.4% shared genes; *r*_*s*_ = 1.00, *p* < 0.01; Figure S7f), and the number of variable sites (798 / 834; 95.7% shared genes; *r*_*s*_ = 0.99, *p* < 0.01; Figure S7i). The other two criteria showed greater differences between NT and AA data matrices (average bootstrap value: 667 / 834; 80.0% shared genes; *r*_*s*_ = 0.78, *p* < 0.01; Figure S7l; treeness / RCV: 644 / 834; 77.2% shared genes; *r*_*s*_ = 0.72, *p* < 0.01; Figure S7o).

### A genome-scale phylogeny for the family Aspergillaceae

NT and AA phylogenomic analyses of the full data matrix and the five subsampled data matrices under three analytical schemes recovered a broadly consistent set of relationships (Figure 1, 2, 3, 4). Across all 36 species-level phylogenies, we observed high levels of topological similarity (average topological similarity: 97.2 ± 2.5%; minimum: 92.2%; maximum: 100%) (Figure 2), with both major genera (*Aspergillus* and *Penicillium*) as well as all sections in *Aspergillus* and *Penicillium* (Houbraken and Samson 2011; Kocsubé et al. 2016) recovered as monophyletic (Figures 1, 3, and 4). Additionally, all but one internodes exhibited absolute UFBoot scores (Hoang et al. 2018); the sole exception was internode 33 (I33), which received 95 UFBoot support (Figure 1 and S9).

**Figure 2.**
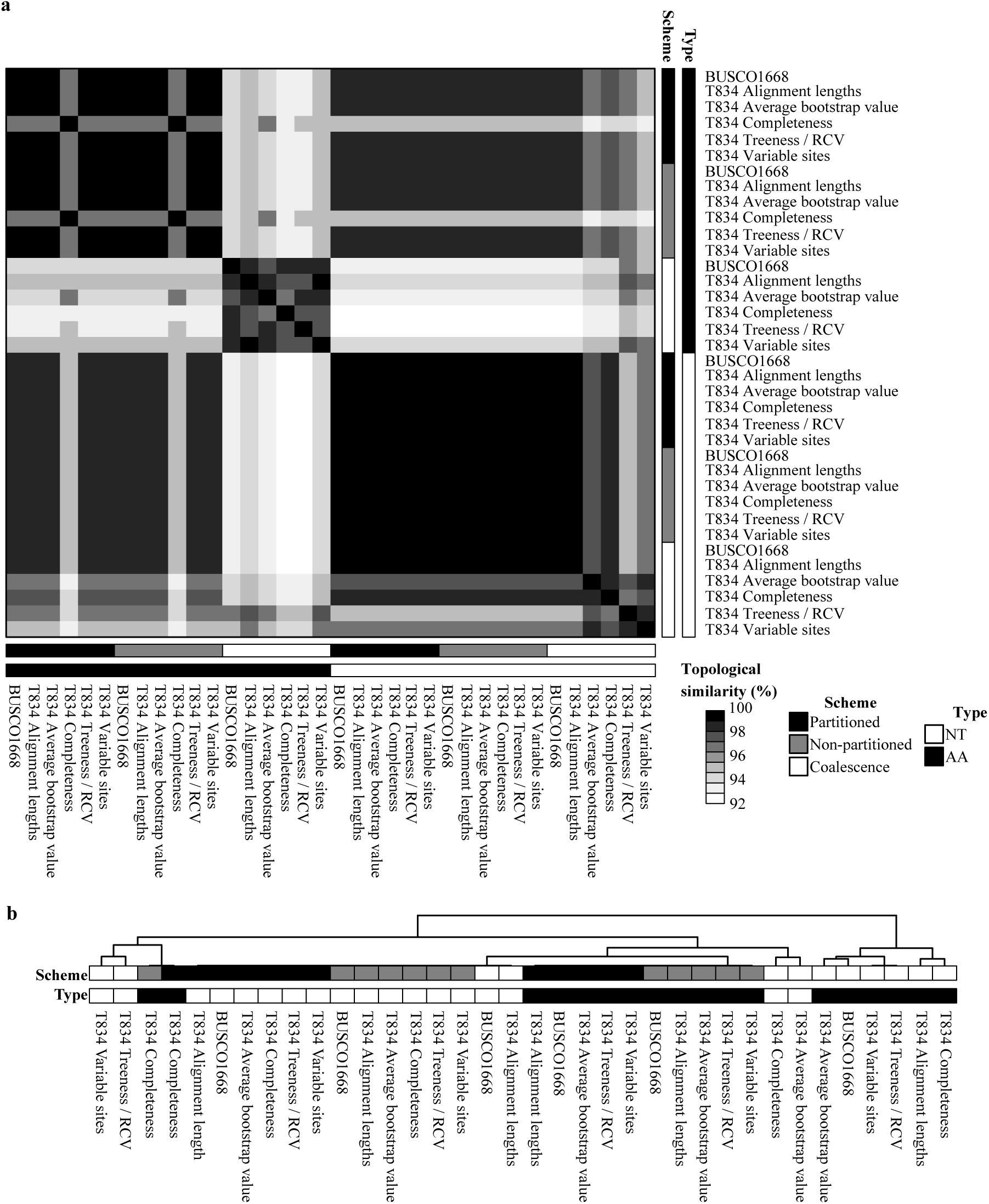
Topological similarity between the 36 phylogenies constructed using 6 different data matrices, 2 different sequence types, and 3 analytical schemes. (a) A heatmap depiction of topological similarity between the 36 phylogenies constructed in this study. The 36 phylogenies were inferred from analyses of 2 different sequence types (i.e., protein: depicted in black; nucleotide: depicted in white), 3 different analytical schemes (i.e., partitioned: depicted in black; non-partitioned: depicted in grey; coalescence: depicted in white), and 6 different matrices (full data matrix: “BUSCO1668”, and 5 subsampled ones, all starting with “T834”; depending on the subsampling strategy, they are identified as “T834 Alignment lengths”, “T834 Average bootstrap value”, “T834 Completeness”, “T834 Treeness / RCV”, and T834 Variable sites”). (b) Hierarchical clustering based on topological similarity values among the 36 phylogenies.

**Figure 3.**
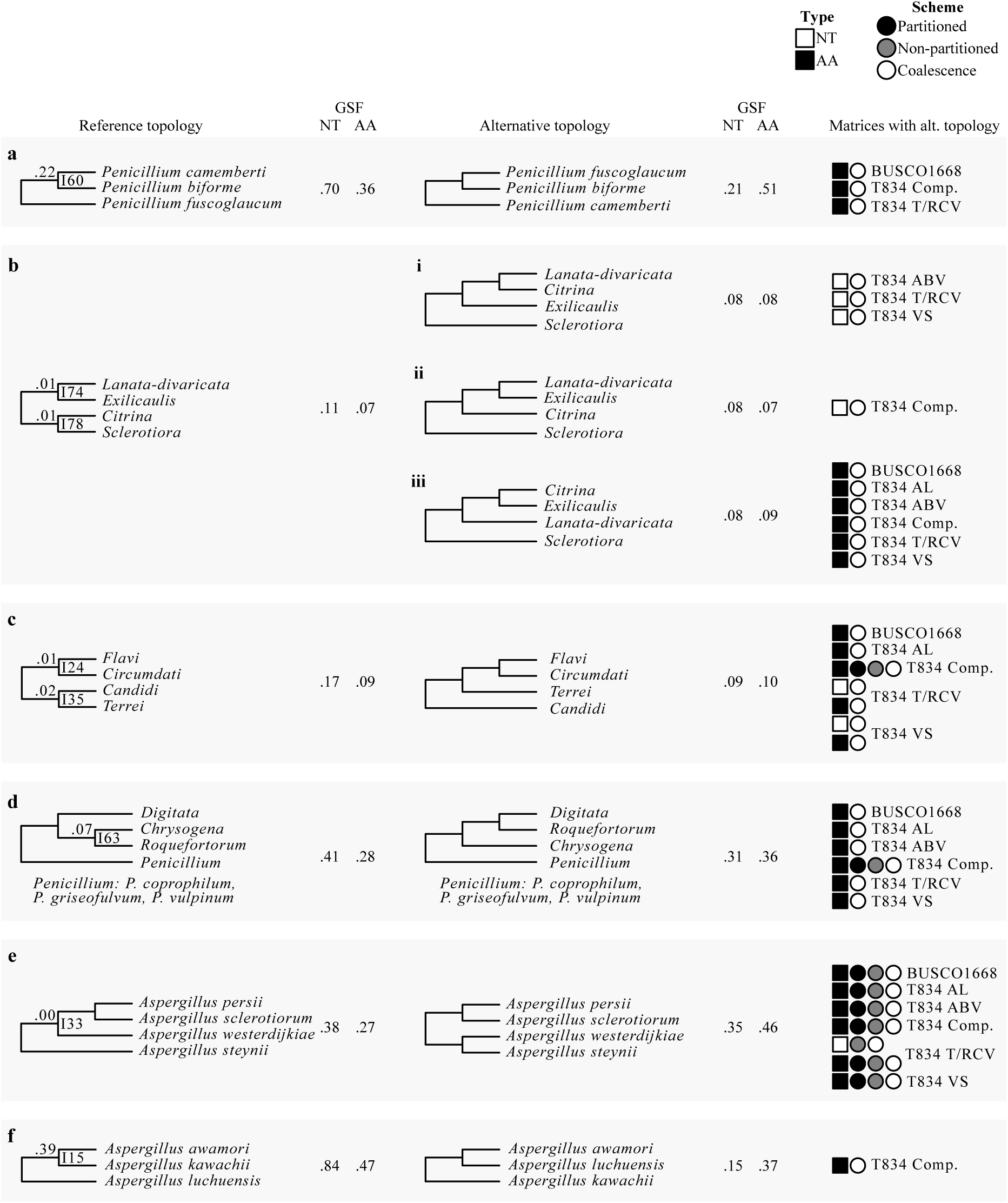
The eight internodes not recovered in all 36 phylogenies. Internode numbers refer to internodes that have at least one conflicting topology among the 36 phylogenetic trees inferred from the full and five subsampled data matrices across three different schemes and two data types. The internode recovered from the analysis of the 1,668-gene nucleotide matrix (Figure 1) is shown on the left and the conflicting internode(s) on the right. Next to each of the internodes, the nucleotide (NT) and amino acid (AA) gene support frequency (GSF) values are shown. On the far right, the sequence type, scheme, and data matrix characteristics of the phylogenies that supports the conflicting internodes are shown. NT and AA sequence types are represented using black and white squares, respectively; partitioned concatenation, non-partitioned concatenation, and coalescence analytical schemes are depicted as black, grey, or white circles, respectively; and the matrix subset is written next to the symbols.

**Figure 4.**
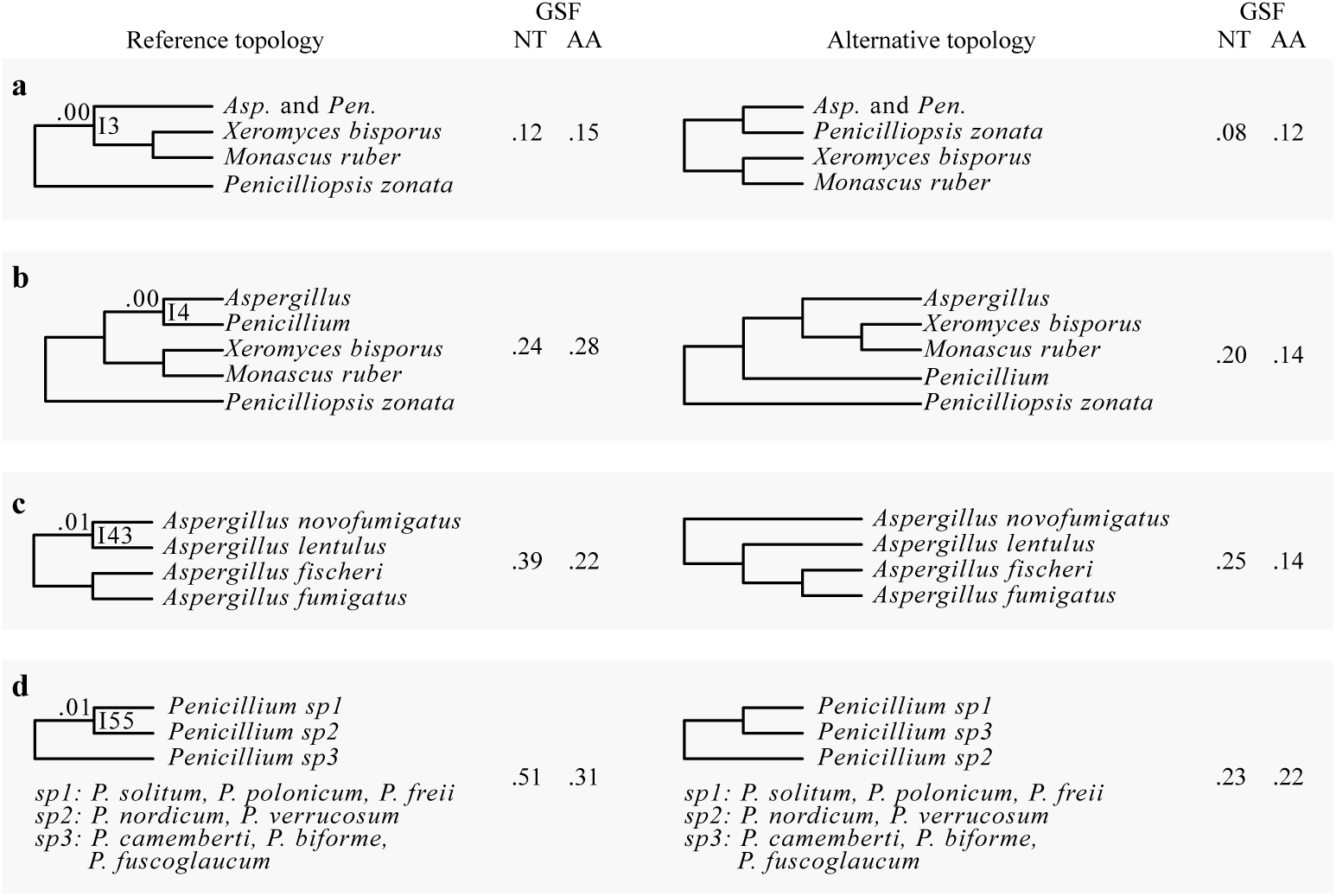
The four internodes recovered in all 36 phylogenies but that exhibit very low internode certainty values. Four bipartitions were recovered by all 36 phylogenies but had internode certainty values below 0.10. The internode recovered from the analysis of all 36 phylogenies, including of the 1,668-gene nucleotide matrix (Figure 1), is shown on the left and the most prevalent, conflicting internode on the right. Next to each of the internodes, the nucleotide (NT) and amino acid (AA) gene support frequency (GSF) values are shown.

Surprisingly, one taxon previously reported to be part of Aspergillaceae, *Basipetospora chlamydospora*, was consistently placed among outgroup species (Figure 1) and may represent a misidentified isolate. To identify the isolate’s true identity, we blasted the nucleotide sequence of *tef1* from the isolate against the “nucleotide collection (nr/nt)” database using MEGABlast (Morgulis et al. 2008) on NCBI’s webserver. We found the top three hits were to *Podospora anserina* (Class Sordariomycetes, PODANS_1_19720; e-value: 0.0, max score: 1753, percent identity: 91%), *Scedosporium apiospermum* (Class Sordariomycetes, SAPIO_CDS5137; e-value: 0.0, max score: 1742, percent identity: 92%), and *Isaria fumosorosea* (Class Sordariomycetes, ISF_05984; e-value: 0.0, max score: 1724, percent identity: 90%). These results make it difficult to ascribe the genome of the misidentified isolate to a specific genus and species but confirm its placement outside of Aspergillaceae; we refer to the isolate by its strain identifier, JCM 23157.

### Examination of the Aspergillaceae phylogeny reveals 12 incongruent bipartitions

Examination of all 36 species-level phylogenies revealed the existence of 8 (8 / 78; 10.3%) incongruent bipartitions. Complementary examination of IC, a bipartition-based measure of incongruence, revealed an additional 4 / 78 (5.1%) bipartitions that displayed very high levels of incongruence at the gene level, raising the total number of incongruent bipartitions to 12 (12 / 78; 15.4%).

Examination of the eight conflicting bipartitions stemming from the comparison of the 36 phylogenies showed that they were very often associated with data type (NT or AA) and scheme employed (concatenation or coalescence). For example, the first instance of incongruence concerns the identity of the sister species to *Penicillium biforme* (I60; Figure 1 and 3a); this species is *P. camemberti* in the reference phylogeny but analyses of the full and two subsampled AA data matrices with coalescence recover instead *Penicillium fuscoglaucum*. The data type and analytical scheme employed also appear to underlie the second and third instances of incongruence, which concern the placement of sections *Exilicaulis* and *Sclerotiora* (I74 and I78; Figures 1 and 3b), the fourth and fifth instances, which concern relationships among *Aspergillus* sections (I24 and I35; Figures 1 and 3c), as well as the sixth instance, which concerns relationships among the sections *Digitata*, *Chrysogena*, and *Roquefortorum* (I63; Figure 1 and 3d). The seventh instance is also associated with data type, but not with the scheme employed; while the reference as well as most subsampled NT matrices support the *Aspergillus persii* and *Aspergillus sclerotiorum* clade as sister to *Aspergillus westerdijkiae* (I33; Figure 1 and 3e), most AA data matrices recover a conflicting bipartition where *A. steynii* is the sister group of *A. westerdijkiae.* The final instance of incongruence was the least well supported, as 35 / 36 (97.2%) phylogenies supported *Aspergillus kawachii* as the sister group to *Aspergillus awamori* (I15, Figure 1 and 3f), but analysis of one AA subsampled data matrix with coalescence instead recovered *Aspergillus luchuensis* as the sister group.

For each of these bipartitions (Figure 3), we examined clustering patterns using multiple correspondence analysis of matrix features (i.e., sequence type and subsampling method) and analysis scheme among trees that support the reference and alternative topologies (Figure S10). Distinct clustering patterns were observed for I74, I78, and I33 (Figure 3 and S10). For I74 and I78, there are three alternative, conflicting topologies, with the first two clustering separately from the third (Figure 3b and S10b). For I33, phylogenies that support the reference and alternative topologies formed distinct clusters (Figure 3e). Examination of the contribution of variables along the second dimension, which is the one that differentiated variables that supported each topology, revealed that the distinct clustering patterns were driven by sequence type (Figure S10g and h).

Examination of IC values revealed four additional bipartitions with strong signatures for incongruence at the gene level, defined as IC score lower than 0.10. The first instance concerns the sister taxon to the *Aspergillus* and *Penicillium* clade. Although all 36 phylogenies recover a clade comprised of *Xeromyces bisporus* and *Monascus ruber* as the sister group, the IC score for this bipartition is 0.00 (I3; Figure 4a); the most prevalent, conflicting bipartition supports *Penicilliopsis zonata* as sister to *Aspergillus* and *Penicillium* (Figure 4a). Similarly, although all 36 phylogenies recover *Penicillium* as sister to *Aspergillus*, the IC score for this bipartition is also 0.00 (I4; Figure 4b); the most prevalent, conflicting bipartition supports *X. bisporus* and *M. ruber* as the sister clade to *Aspergillus* (Figure 4b). In the third instance, all 36 phylogenies support *Aspergillus novofumigatus* and *Aspergillus lentulus* as sister species, but the IC score of this bipartition is 0.01 (I43; Figure 4c); the most prevalent, conflicting bipartition recovers *A. lentulus* as the sister species to a clade comprised of *Aspergillus fumigatus* and *Aspergillus fischeri* (Figure 4c). Finally, all 36 phylogenies supported a clade of *Penicillium solitum, Penicillium polonicum*, and *Penicillium freii* as sister to a clade of *Penicillium nordicum* and *Penicillium verrucosum*, but the IC score for this bipartition is 0.01 (I55; Figure 4d); the most prevalent, conflicting bipartition supports the clade of *P. solitum, P. polonicum,* and *P. freii* as sister to a clade of *P. camemberti, P. biforme* and *P. fuscoglaucum* (Figure 4d).

To examine the underlying individual gene support to the resolution of these 12 bipartitions, we examined the phylogenetic signal contributed by each individual gene in the full NT data matrix. In all 12 bipartitions, we found that inferences were robust to single gene outliers with strong phylogenetic signal (Figure S11; File S4).

### Incongruence in the Aspergillaceae phylogeny

Examination of the 12 incongruent bipartitions with respect to their placement on the phylogeny (shallow, i.e., near the tips of the phylogeny or deeper, i.e., away from the tips and toward the base of the phylogeny) and the amount of conflict (quantified using IC and GSF) allowed us to group them into three categories: (i) shallow bipartitions (I15 and I60) with low levels of incongruence, (ii) shallow bipartitions (I33, I43, and I55) with high levels of incongruence, and (iii) deeper bipartitions (I3, I4, I24, I35, I63, I74, and I78) with varying levels of incongruence and typically associated with single taxon long branches.

#### (i) Shallow bipartitions with low levels of incongruence

The two bipartitions that fell into this category, I60 (Figure 3a) and I15 (Figure 3f), exhibited low levels of incongruence among closely related taxa. For I60, the reference bipartition was observed in 33 / 36 phylogenies, had an IC score of 0.22, and GSF_NT_ and GSF_AA_ scores of 0.70 and 0.21, respectively. Similarly, the reference bipartition for I15 was observed in 35 / 36 phylogenies, had an IC score of 0.39, and GSF_NT_ and GSF_AA_ scores of 0.84 and 0.47, respectively. Notably, the GSF_NT_ scores were substantially higher for the reference bipartitions in both of these cases.

#### (ii) Shallow bipartitions with high levels of incongruence

The three shallow bipartitions, I33 (Figure 3e), I43 (Figure 4c), and I55 (Figure 4d), in this category exhibited high levels of incongruence among closely related taxa. For I33, the reference bipartition was observed in 16 / 36 (44.4%), had an IC score of 0.00, and GSF_NT_ and GSF_AA_ scores of 0.38 and 0.27, respectively. The reference bipartition for I43 was observed in all 36 phylogenies, had an IC score of 0.01 and GSF_NT_ and GSF_AA_ scores of 0.39 and 0.22, respectively. Similarly, the reference bipartition I55 was observed in all 36 phylogenies, had an IC score of 0.01, and GSF_NT_ and GSF_AA_ scores of 0.51 and 0.31, respectively. Notably, in all three cases, substantial fractions of genes supported both the reference and the conflicting bipartitions, with both the GSF_NT_ and GSF_AA_ scores of each pair of bipartitions being almost always higher than 0.2.

#### (iii) Deeper bipartitions often associated with single taxon long branches

The seven bipartitions in this category were I74 and I78 (Figure 3b), I24 and I35 (Figure 3c), I63 (Figure 3d), I3 (Figure 4a), and I4 (Figure 4b). All of them are located deeper in the tree and most involve single taxa with long terminal branches (Figure 1). The reference bipartitions for internodes I74 and I78, which concern relationships among the sections *Lanata-divaricata*, *Exilicaulis, Citrina*, and *Sclerotiora* were observed in 26 / 36 (72.2%) phylogenies; the remaining 10 / 36 (27.8%) phylogenies recovered three alternative, conflicting bipartitions. Both reference bipartitions had IC scores of 0.01, and GSF_NT_ and GSF_AA_ scores of 0.11 and 0.07, respectively. The reference bipartitions for internodes I24 and I35, which concern the placement of *Aspergillus terreus*, the single taxon representative of section *Terrei*, were observed in 27 / 36 (75.0%) phylogenies, had IC scores of 0.01 and 0.02, and GSF_NT_ and GSF_AA_ scores of 0.17 and 0.09, respectively. The reference bipartition I63, which involved the placement of the *Penicillium digitatum*, the sole representative of section *Digitata*, was observed in 28 / 36 (77.8%), had an IC score of 0.07, and GSF_NT_ and GSF_AA_ scores of 0.41 and 0.28, respectively. Finally, the reference bipartitions I3 and I4 (Figure 4), which concern the identity of the sister taxon of *Aspergillus* and *Penicillium* (I3) and the identity of the sister taxon of *Aspergillus* (I4), were not observed among the 36 phylogenies but both had IC values of 0.00. For I3, GSF_NT_ and GSF_AA_ scores were 0.12 and 0.15, respectively. For I4, GSF_NT_ and GSF_AA_ scores were 0.24 and 0.28, respectively.

### Topology tests

The phylogeny of the genera *Aspergillus* and *Penicillium* has been a topic of debate. Our topology supports the reciprocal monophyly of *Aspergillus* and *Penicillium* and rejects the monophyly of narrow *Aspergillus*. Both of these results are consistent with some previous studies (Kocsubé et al. 2016) (Figure 6) but in contrast to other previous studies, which recovered a topology where *Penicillium* is sister to section *Nidulantes* within *Aspergillus* and narrow *Aspergillus* (sections *Nigri*, *Ochraceorosei*, *Flavi*, *Circumdati*, *Candidi*, *Terrei*) was monophyletic (Pitt and Taylor 2014; Taylor et al. 2016). To further evaluate both of these hypotheses, we conducted separate topology constraint analyses using the Shimodaira-Hasegawa (Shimodaira and Hasegawa 1999) and the approximately unbiased tests (Shimodaira 2002). Both tests rejected the constrained topologies (Table 1; p-value < 0.001 for all tests), providing further support that *Aspergillus* and *Penicillium* are reciprocally monophyletic and that narrow *Aspergillus* is not monophyletic (Figure 6).

**Table 1.**
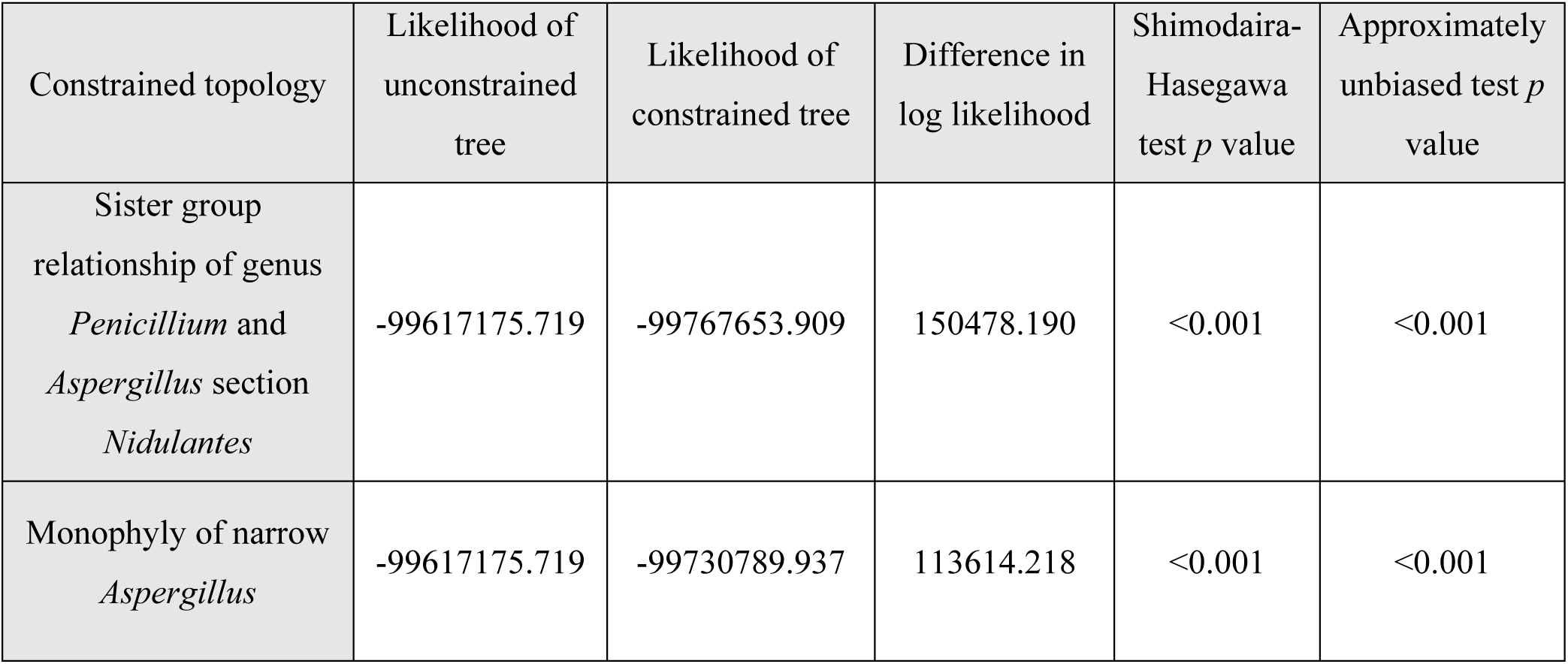
Topology tests reject the sister group relationship of genus *Penicillium* and *Aspergillus* section *Nidulantes* as well as the monophyly of narrow *Aspergillus.*

| Constrained topology | Likelihood of unconstrained tree | Likelihood of constrained tree | Difference in log likelihood | Shimodaira-Hasegawa test *p* value | Approximately unbiased test *p* value |
| --- | --- | --- | --- | --- | --- |
| Sister group relationship of genus *Penicillium* and *Aspergillus* section *Nidulantes* | -99617175.719 | -99767653.909 | 150478.190 | <0.001 | <0.001 |
| Monophyly of narrow *Aspergillus* | -99617175.719 | -99730789.937 | 113614.218 | <0.001 | <0.001 |

### A geological timeline for the evolutionary diversification of the Aspergillaceae family

To estimate the evolutionary diversification among *Aspergillaceae*, we subsampled the 1,668-gene matrix for high-quality genes with “clock-like” rates of evolution by examining DVMC (Liu et al. 2017) values among single gene trees. Examination of the DVMC values facilitated the identification of a tractable set of high-quality genes for relaxed molecular clock analyses (Figure S12). We found that *Aspergillaceae* originated 117.4 (95% CI: 141.5 - 96.9) mya during the Cretaceous period (Figure 5). We found that the common ancestor of *Aspergillus* and *Penicillium* split from the *X. bisporus* and *M. ruber* clade shortly thereafter, approximately 109.8 (95% CI: 129.3 - 93.5) mya. We also found that the genera *Aspergillus* and *Penicillium* split 94.0 (95% CI: 106.8 - 83.0) mya, with the last common ancestor of *Aspergillus* originating approximately 81.7 mya (95% CI: 87.5 - 72.9) and the last common ancestor of *Penicillium* originating approximately 73.6 mya (95% CI: 84.8 - 60.7).

**Figure 5.**
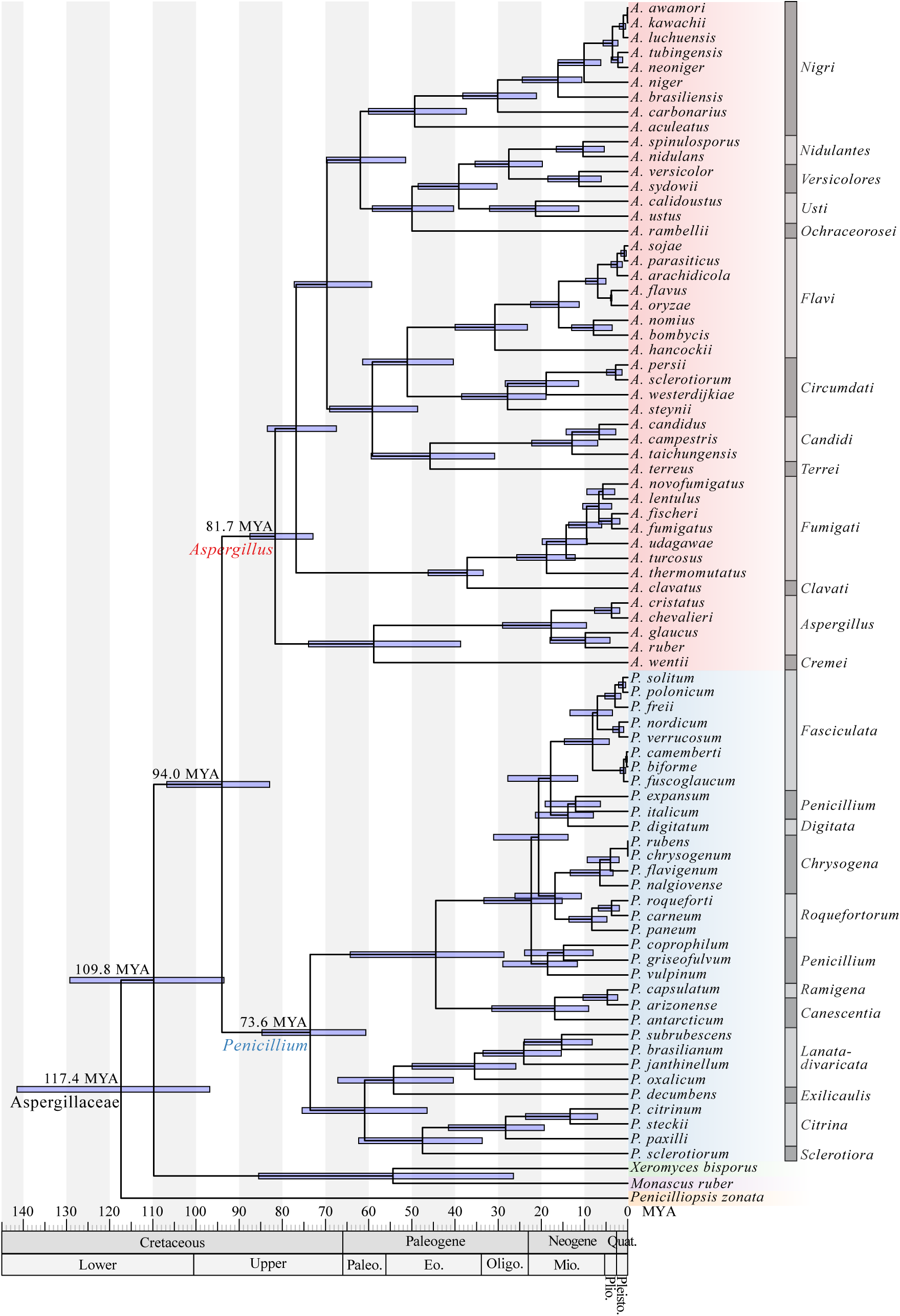
A molecular timetree for the family Aspergillaceae. Blue boxes around each internode correspond to 95% divergence time confidence intervals for each branch of the Aspergillaceae phylogeny. For reference, the geologic time scale is shown right below the phylogeny. Different genera are depicted using different colored boxes; *Aspergillus* is shown in red, *Penicillium* in blue, *Xeromyces* in green, *Monascus* in purple, and *Penicilliopsis* in orange. Different sections within *Aspergillus* and *Penicillium* are depicted with alternating dark grey and grey bars. Dating estimates were calibrated using the following constraints: origin of Aspergillaceae (I2; 50-146 million years ago [mya]), origin of *Aspergillus* (I5; 43-85 mya) the *A. flavus* and *A. oryzae* split (I30; 3.68-3.99 mya), and the *A. fumigatus* and *A. clavatus* split (I38; 35-39 mya); all constraints were obtained from TIMETREE (Hedges et al. 2006).

Among *Aspergillus* sections, section *Nigri*, which includes the industrial workhorse *A. niger*, originated 49.4 (95% CI: 60.1 - 37.4) mya. Section *Flavi*, which includes the food fermenters *A. oryzae* and *A. sojae* and the toxin-producing, post-harvest food contaminant, and opportunistic animal and plant pathogen *A. flavus*, originated 30.8 (95% CI: 40.0 - 23.3) mya. Additionally, section *Fumigati*, which includes the opportunistic human pathogen *A. fumigatus*, originated 18.8 (95% CI: 25.7 - 12.2) mya. Among *Penicillium* sections, section *Fasciculata*, which contains Camembert and Brie cheese producer *P. camemberti* and the ochratoxin A producer, *P. verrucosum*, originated 8.1 (95% CI: 14.7 - 4.3) mya. Section *Chrysogena,* which includes the antibiotic penicillin producing species *P. chrysogenum*, originated 6.5 (95% CI: 13.3 - 3.4) mya. Additionally, section *Citrina*, which contains *P. citrinum*, which the first statin was isolated from and is commonly associated with moldy citrus fruits (Endo et al. 1976), originated 28.3 (95% CI: 41.5 - 19.3) mya.

Finally, our analysis also provides estimates of the origins of various iconic pairs of species within *Aspergillus* and *Penicillium*. For example, among *Aspergillus* species pairs, we estimate that *A. fumigatus* and the closest relative with a sequenced genome, *A. fischeri* (Mead et al. 2018), diverged 3.7 (95% CI: 6.7 – 1.9) mya and *Aspergillus flavus* and the domesticated counterpart, *A. oryzae* (Gibbons et al. 2012), 3.8 (95% CI: 4.0 – 3.7) mya. Among *Penicillium* species pairs, we estimate *P. camemberti*, which contributes to cheese production to have diverged from its sister species and cheese contaminant *P. biforme* (Ropars et al. 2012) approximately 0.3 (95% CI: 0.5 – 0.1) mya. Finally, we estimate that *P. roqueforti*, another species that contributes to cheese production, diverged from its close relative *P. carneum* (Ropars et al. 2012) 3.8 (95% CI: 6.8 – 2.0) mya.

## Discussion

Our analyses provide a robust evaluation of the evolutionary relationships and diversification among Aspergillaceae, a family of biotechnologically and medically significant fungi. We scrutinized our proposed reference phylogeny (Figure 1) against 35 other phylogenies recovered using all possible combinations of six multi-gene data matrices (full or subsamples thereof), three maximum likelihood schemes, and two sequence types and complemented this analysis with bi-partitioned based measures of support (Figures 1 and 2). Through these analyses, we found that 12 / 78 (15.4%) bipartitions were incongruent (Figure 3 and 4) and explored the characteristics as well as sources of these instances of incongruence. Finally, we placed the evolution and diversification of Aspergillaceae in the context of geological time.

Comparison of our 81-taxon, 1,668-gene phylogeny to a previous one based on a maximum likelihood analysis of 9 loci for 204 Aspergillaceae species (Kocsubé et al. 2016), suggests that our analyses identified and strongly supported several new relationships and resolved previously poorly supported bipartitions (Figure 1, Figure 6). The robust resolution of our phylogeny is likely due to the very large size of our data matrix, both in terms of genes as well as in terms of taxa. For example, the placement of *Aspergillus* section *Nigri* has been unstable in previous phylogenomic analyses (Figure S1) (Yang et al. 2016; de Vries et al. 2017; Kjærbølling et al. 2018), but our denser sampling of taxa in this section as well as inclusion of representative taxa from sections *Nidulantes, Versicolores, Usti*, and *Ochraceorosei* now provides strong support for the sister relationship of the *Aspergillus* section *Nigri* to sections *Nidulantes, Versicolores, Usti*, and *Ochraceorosei* (Figure 1).

**Figure 6.**
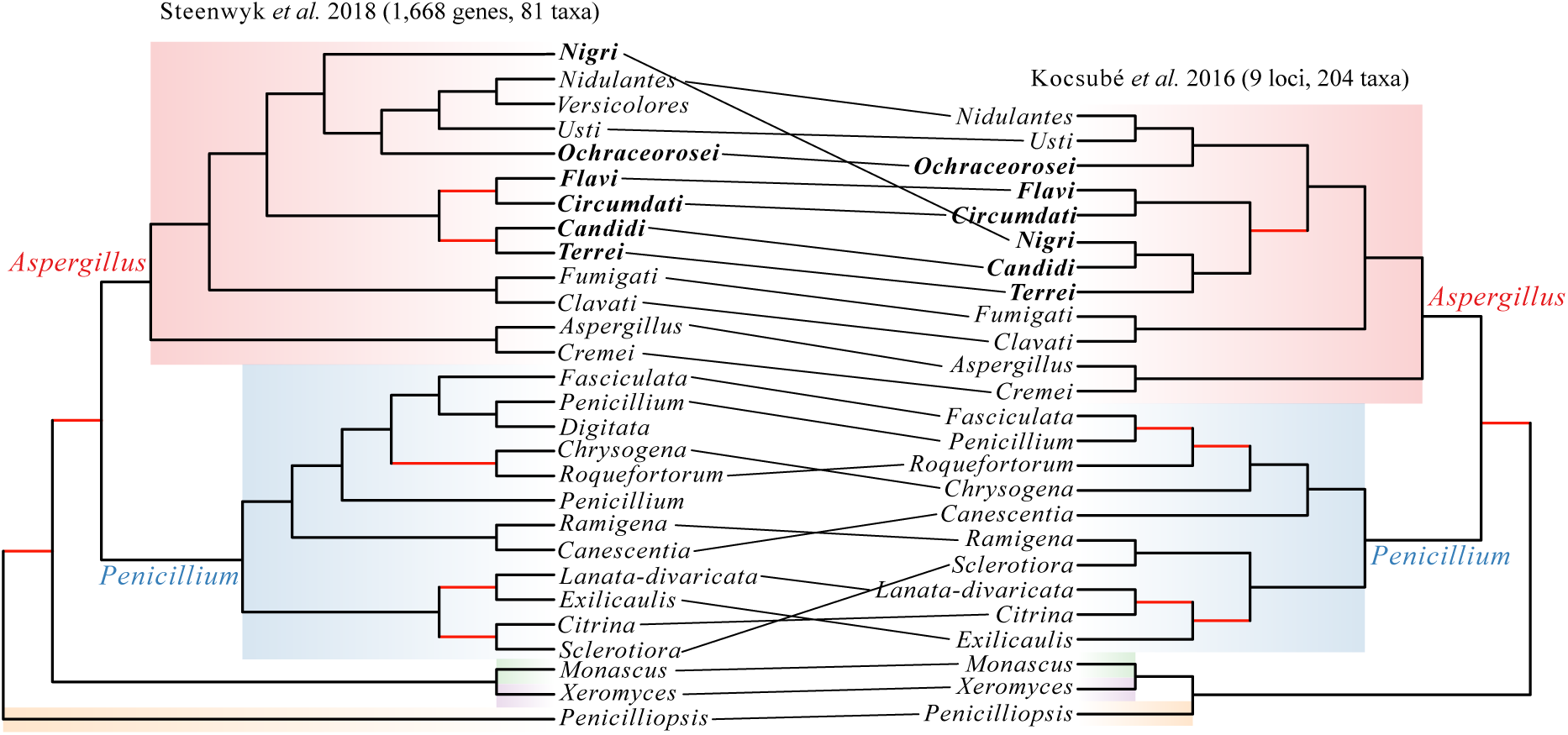
A visual comparison of the differences between the phylogeny reported in this study and the phylogeny reported in Kocsubé *et al.* 2016. Tanglegram between the section-level phylogeny presented in this study (left) and the section-level phylogeny presented by Kocsubé *et al*. 2016 (right). The key differences between the two phylogenies lie in the placements of sections *Nigri*, *Ramigena*, and *Canescentia*. Species in bold belong to narrow *Aspergillus* and red branches represent bipartitions that are not robustly supported in each study.

However, our analysis also identified several relationships that exhibit high levels of incongruence (Figures 3 and 4). In general, gene tree incongruence can stem from biological or analytical factors (Rokas et al. 2003; Shen et al. 2017). Biological processes such as incomplete lineage-sorting (ILS) (Degnan and Salter 2005), hybridization (Sang and Zhong 2000), gene duplication and subsequent loss (Hallett et al. 2004), horizontal gene transfer (Doolittle and Bapteste 2007), and natural selection (Castoe et al. 2009; Li et al. 2010), can cause the histories of genes to differ from one another and from the species phylogeny. Importantly, although the expected patterns of incongruence will be different for each factor and depend on a number of parameters, the observed patterns of conflict in each of the 12 cases of incongruence in the Aspergillaceae phylogeny can yield insights and allow the formation of hypotheses about the potential drivers in each case. For example, ILS often results in relatively low levels of incongruence; for instance, examination of the human, chimp, and gorilla genomes has showed that 20-25% of the gene histories differ from the species phylogeny (Patterson et al. 2006; Hobolth et al. 2007). In contrast, recent hybridization is expected to typically produce much higher levels of incongruence due to rampant sequence similarity among large amounts of genomic content; for instance, examination of *Heliconius* butterfly genomes revealed incongruence levels higher than 40% (Martin et al. 2013).

Additionally, analytical factors such as model choice (Phillips et al. 2004) and taxon sampling (Rokas and Carroll 2005; Nabhan and Sarkar 2012) can lead to erroneous inference of gene histories. Perhaps the most well-known instance of incongruence stemming from analytical factors is what is known as “long branch attraction”, namely the situation where highly divergent taxa, i.e., the ones with the longest branches in the phylogeny, will often artifactually group with other long branches (Baldauf 2003).

Examination of the patterns of incongruence in the Aspergillaceae phylogeny allows us to not only group the 12 incongruent internodes with respect to their patterns of conflict but also to postulate putative drivers of the observed incongruence. For example, both I15 and I60 are shallow internodes exhibiting low levels of incongruence, suggesting that one likely driver of the observed incongruence is ILS. In contrast, the shallow internodes I33, I43, and I55 exhibit much higher levels of incongruence that are most likely to be the end result of processes, such as hybridization or repeated introgression. Finally, the remaining seven incongruent internodes (I3, I4, I24, I35, I63, I74, and I78) exhibit varying levels of incongruence and are typically associated with single taxon long branches (Figures 1, 3, and 4), implicating taxon sampling as a likely driver of the observed incongruence. Given that inclusion of additional taxa robustly resolved the previously ambiguous placement of the long-branched *Aspergillus* section *Nigri* (see discussion above), we predict that additional sampling of taxa that break up the long branches associated with these seven internodes will lead to their robust resolution.

Notably, the topology of our phylogeny was able to resolve two contentious issues that emerged from analyses of data matrices containing a few genes (Kocsubé et al. 2016; Taylor et al. 2016) and that are important for taxonomic relationships within the family. Specifically, our phylogenetic analyses rejected the sister group relationship of genus *Penicillium* and *Aspergillus* section *Nidulantes* as well as the monophyly of a group of *Aspergillus* sections that are referred to as narrow *Aspergillus* (Table 1, p-value < 0.001 for all tests). Instead, our phylogeny shows that the genera *Aspergillus* and *Penicillium* are reciprocally monophyletic. These results are consistent with the current nomenclature proposed by the International Commission of *Penicillium* and *Aspergillus* (https://www.aspergilluspenicillium.org/), and inconsistent with the phylogenetic arguments put forward in proposals for taxonomic revision (Taylor et al. 2016). However, it should be noted that our study did not include representatives of the genera *Phialosimplex* and *Polypaecilum*, which lack known asexual stages, and appear to be placed within the genus *Aspergillus* (Kocsubé et al. 2016; Taylor et al. 2016). *Basipetospora* species also lack known asexual stages and are also placed within *Aspergillus* (Kocsubé et al. 2016; Taylor et al. 2016); unfortunately, the sole genome sequenced from this genus, JCM 23157, appears to be a contaminant from the class Sordariomycetes (Figure 1).

Finally, our relaxed molecular clock analysis of the Aspergillaceae phylogeny provides a robust and comprehensive time-scale for the evolution of Aspergillaceae and its two large genera, *Aspergillus* and *Penicillium* (Figure 5), filling a gap in the literature. Previous molecular clock studies provided estimates for only four internodes, mostly within the genus *Aspergillus* (Berbee and Taylor 2001; Hedges et al. 2006; Vijaykrishna et al. 2006; Kensche et al. 2008; Sharpton et al. 2009; Da Lage et al. 2013; Beimforde et al. 2014; Fan et al. 2015; Gaya et al. 2015) and yielded much broader time intervals. For example, the previous estimate for the origin of Aspergillaceae spanned nearly 100 mya (50-146 mya: Berbee and Taylor 2001; Vijaykrishna et al. 2006; Sharpton et al. 2009) while our dataset and analysis provided a much narrower range of 44.7 mya (mean: 117.4; 95% CI: 141.5 - 96.9). Notably, the estimated origins of genera *Aspergillus* (~81.7 mya) and *Penicillium* (~73.6 mya) appear to be comparable to those of other well-known filamentous fungal genera, such as *Fusarium*, whose date of origin has been estimated at ~91.3 mya (Ma et al. 2013; O’Donnell et al. 2013).

## Conclusion

Fungi from Aspergillaceae have diverse ecologies and play significant roles in biotechnology and medicine. Although most of the 81 genomes from Aspergillaceae are skewed towards two iconic genera, *Aspergillus* and *Penicillium*, and do not fully reflect the diversity of the family, they do provide a unique opportunity to examine the evolutionary history of these important fungi using a phylogenomic approach. Our scrutiny of the Aspergillaceae phylogeny, from the Cretaceous to the present, provides strong support for most relationships within the family as well as identifies a few that deserve further examination. Our results suggest that the observed incongruence is likely associated with diverse processes such as incomplete lineage sorting, hybridization and introgression, as well as with analytical issues associated with poor taxon sampling. Our elucidation of the tempo and pattern of the evolutionary history of Aspergillaceae aids efforts to develop a robust taxonomic nomenclature for the family and provides a robust phylogenetic and temporal framework for investigation the evolution of pathogenesis, secondary metabolism, and ecology of this diverse and important fungal family.

## Data availability

All data matrices, species-level and single-gene phylogenies will be available through the figshare repository upon acceptance for publication. The genome sequence and raw reads of *Aspergillus spinulosporus* have been uploaded to GenBank as BioProject PRJNA481010.

## Acknowledgements

We thank members of the Rokas laboratory for helpful suggestions and discussion, Nathan McDonald for technical help with DNA extraction, Mikael Andersen for help with the taxonomic identity of one of the *Aspergillus* genomes included in our study, Joseph Walker and Xiaofan Zhou for useful feedback on the topology constraint analyses, and David Geiser and an anonymous reviewer for constructive comments on the manuscript. This work was conducted, in part, using the Advanced Computing Center for Research and Education at Vanderbilt University. J.L.S. was supported by the Graduate Program in Biological Sciences at Vanderbilt University. This work was supported, in part, by the National Science Foundation (DEB-1442113 to A.R.), the Burroughs Wellcome Fund, and a Guggenheim fellowship (to A.R.).

## References

Albert AW, Chen J, Kuron G, Hunt V, Huff J, Hoffman C, Rothrock J, Lopez M, Joshua H, Harris E, et al. 1980. Mevinolin: a highly potent competitive inhibitor of hydroxymethylglutaryl-coenzyme A reductase and a cholesterol-lowering agent. Proc. Natl. Acad. Sci. U. S. A. 77:3957–3961.

Aminov RI. 2010. A Brief History of the Antibiotic Era: Lessons Learned and Challenges for the Future. Front. Microbiol. 1:134.

Arcila D, Ortí G, Vari R, Armbruster JW, Stiassny MLJ, Ko KD, Sabaj MH, Lundberg J, Revell LJ, Betancur-R. R. 2017. Genome-wide interrogation advances resolution of recalcitrant groups in the tree of life. Nat. Ecol. Evol. 1:0020.

Baldauf SL. 2003. Phylogeny for the faint of heart: a tutorial. Trends Genet. 19:345–351.

Ballester A-R, Marcet-Houben M, Levin E, Sela N, Selma-Lázaro C, Carmona L, Wisniewski M, Droby S, González-Candelas L, Gabaldón T. 2015. Genome, Transcriptome, and Functional Analyses of *Penicillium expansum* Provide New Insights Into Secondary Metabolism and Pathogenicity. Mol. Plant-Microbe Interact. 28:232–248.

Bankevich A, Nurk S, Antipov D, Gurevich AA, Dvorkin M, Kulikov AS, Lesin VM, Nikolenko SI, Pham S, Prjibelski AD, et al. 2012. SPAdes: A New Genome Assembly Algorithm and Its Applications to Single-Cell Sequencing. J. Comput. Biol. 19:455–477.

Beimforde C, Feldberg K, Nylinder S, Rikkinen J, Tuovila H, Dörfelt H, Gube M, Jackson DJ, Reitner J, Seyfullah LJ, et al. 2014. Estimating the Phanerozoic history of the Ascomycota lineages: Combining fossil and molecular data. Mol. Phylogenet. Evol. 78:386–398.

Bell MA, Lloyd GT. 2015. strap: an R package for plotting phylogenies against stratigraphy and assessing their stratigraphic congruence.Smith A, editor. Palaeontology 58:379–389.

Benson DA, Karsch-Mizrachi I, Lipman DJ, Ostell J, Wheeler DL. 2007. GenBank. Nucleic Acids Res. 36:D25–D30.

Berbee ML, Taylor JW. 2001. Fungal molecular evolution: gene trees and geologic time. In: The Mycota. p. 229–246.

Camacho C, Coulouris G, Avagyan V, Ma N, Papadopoulos J, Bealer K, Madden TL. 2009. BLAST+: architecture and applications. BMC Bioinformatics 10:421.

Capella-Gutierrez S, Silla-Martinez JM, Gabaldon T. 2009. trimAl: a tool for automated alignment trimming in large-scale phylogenetic analyses. Bioinformatics 25:1972–1973.

Castoe TA, de Koning APJ, Kim H-M, Gu W, Noonan BP, Naylor G, Jiang ZJ, Parkinson CL, Pollock DD. 2009. Evidence for an ancient adaptive episode of convergent molecular evolution. Proc. Natl. Acad. Sci. 106:8986–8991.

Chain E, Florey HW, Gardner AD, Heatley NG, Jennings MA, Orr-Ewing J, Sanders AG. 1940. Pencillin as a chemotherapeutic agent. Lancet 236:226–228.

Chernomor O, von Haeseler A, Minh BQ. 2016. Terrace Aware Data Structure for Phylogenomic Inference from Supermatrices. Syst. Biol. 65:997–1008.

Chikhi R, Medvedev P. 2014. Informed and automated k-mer size selection for genome assembly. Bioinformatics 30:31–37.

Cock PJA, Antao T, Chang JT, Chapman BA, Cox CJ, Dalke A, Friedberg I, Hamelryck T, Kauff F, Wilczynski B, et al. 2009. Biopython: freely available Python tools for computational molecular biology and bioinformatics. Bioinformatics 25:1422–1423.

Conway JR, Lex A, Gehlenborg N. 2017. UpSetR: an R package for the visualization of intersecting sets and their properties.Hancock J, editor. Bioinformatics 33:2938–2940.

Degnan JH, Salter LA. 2005. Gene tree distributions under the coalescent process. Evolution (N. Y). 59:24–37.

Doolittle WF, Bapteste E. 2007. Pattern pluralism and the Tree of Life hypothesis. Proc. Natl. Acad. Sci. 104:2043–2049.

Eddy SR. 2011. Accelerated Profile HMM Searches.Pearson WR, editor. PLoS Comput. Biol. 7:e1002195.

Edwards SV. 2009. Is a new and general theory of molecular systematics emerging? Evolution (N. Y). 63:1–19.

Endo A. 2010. A historical perspective on the discovery of statins. Proc. Japan Acad. Ser. B 86:484–493.

Endo A, Kuroda M, Tsujita Y. 1976. ML-236A, ML-236B, and ML-236C, new inhibitors of cholesterogensis produced by *Penicillium citrinum*. J. Antibiot. (Tokyo). 29:1346–1348.

Fan H-W, Noda H, Xie H-Q, Suetsugu Y, Zhu Q-H, Zhang C-X. 2015. Genomic Analysis of an Ascomycete Fungus from the Rice Planthopper Reveals How It Adapts to an Endosymbiotic Lifestyle. Genome Biol. Evol. 7:2623–2634.

Felsenstein J. 1981. Evolutionary trees from DNA sequences: A maximum likelihood approach. J. Mol. Evol. 17:368–376.

Felsenstein J. 1986. The Newick tree format. English.

Felsenstein J. 1996. Inferring phylogenies from protein sequences by parsimony, distance, and likelihood methods. Methods Enzymol. 266:418–427.

Fleming A. 1980. On the Antibacterial Action of Cultures of a *Penicillium*, with Special Reference to Their Use in the Isolation of *B. influenzae*. Clin. Infect. Dis. 2:129–139.

Fox EM, Howlett BJ. 2008. Secondary metabolism: regulation and role in fungal biology. Curr. Opin. Microbiol. 11:481–487.

Frisvad JC, Larsen TO. 2015. Chemodiversity in the genus *Aspergillus*. Appl. Microbiol. Biotechnol. 99:7859–7877.

Gaya E, Fernández-Brime S, Vargas R, Lachlan RF, Gueidan C, Ramírez-Mejía M, Lutzoni F. 2015. The adaptive radiation of lichen-forming Teloschistaceae is associated with sunscreening pigments and a bark-to-rock substrate shift. Proc. Natl. Acad. Sci. 112:11600–11605.

Gibbons JG, Rokas A. 2013. The function and evolution of the *Aspergillus* genome. Trends Microbiol. 21:14–22.

Gibbons JG, Salichos L, Slot JC, Rinker DC, McGary KL, King JG, Klich MA, Tabb DL, McDonald WH, Rokas A. 2012. The Evolutionary Imprint of Domestication on Genome Variation and Function of the Filamentous Fungus *Aspergillus oryzae*. Curr. Biol. 22:1403–1409.

Gurevich A, Saveliev V, Vyahhi N, Tesler G. 2013. QUAST: quality assessment tool for genome assemblies. Bioinformatics 29:1072–1075.

Hallett M, Lagergren J, Tofigh A. 2004. Simultaneous identification of duplications and lateral transfers. In: Proceedings of the eighth annual international conference on Computational molecular biology - RECOMB’04. New York, New York, USA: ACM Press. p. 347–356.

Harrell Jr FE. 2015. Package “Hmisc” (v4.0-0).

Hedayati MT, Pasqualotto AC, Warn PA, Bowyer P, Denning DW. 2007. *Aspergillus flavus*: human pathogen, allergen and mycotoxin producer. Microbiology 153:1677–1692.

Hedges SB, Dudley J, Kumar S. 2006. TimeTree: A public knowledge-base of divergence times among organisms. Bioinformatics 22:2971–2972.

Hess J, Goldman N. 2011. Addressing Inter-Gene Heterogeneity in Maximum Likelihood Phylogenomic Analysis: Yeasts Revisited.Rattray M, editor. PLoS One 6:e22783.

Hoang DT, Chernomor O, von Haeseler A, Minh BQ, Vinh LS. 2018. UFBoot2: Improving the Ultrafast Bootstrap Approximation. Mol. Biol. Evol. 35:518–522.

Hobolth A, Christensen OF, Mailund T, Schierup MH. 2007. Genomic Relationships and Speciation Times of Human, Chimpanzee, and Gorilla Inferred from a Coalescent Hidden Markov Model. PLoS Genet. 3:e7.

Houbraken J, Samson RA. 2011. Phylogeny of *Penicillium* and the segregation of Trichocomaceae into three families. Stud. Mycol. 70:1–51.

Houbraken J, de Vries RP, Samson RA. 2014. Modern Taxonomy of Biotechnologically Important *Aspergillus* and *Penicillium* Species. In: Advances in Applied Microbiology. Vol. 86. p. 199–249.

Ioannidis P, Simao FA, Waterhouse RM, Manni M, Seppey M, Robertson HM, Misof B, Niehuis O, Zdobnov EM. 2017. Genomic features of the damselfly *Calopteryx splendens* representing a sister clade to most insect orders. Genome Biol. Evol. 9:415–430.

Jones DT, Taylor WR, Thornton JM. 1992. The rapid generation of mutation data matrices from protein sequences. Bioinformatics 8:275–282.

Junier T, Zdobnov EM. 2010. The Newick utilities: high-throughput phylogenetic tree processing in the UNIX shell. Bioinformatics 26:1669–1670.

Kalyaanamoorthy S, Minh BQ, Wong TKF, von Haeseler A, Jermiin LS. 2017. ModelFinder: fast model selection for accurate phylogenetic estimates. Nat. Methods 14:587–589.

Kassambara A, Mundt F. 2017. factoextra. R Packag. v. 1.0.5.

Katoh K, Standley DM. 2013. MAFFT Multiple Sequence Alignment Software Version 7: Improvements in Performance and Usability. Mol. Biol. Evol. 30:772–780.

Keller NP, Turner G, Bennett JW. 2005. Fungal secondary metabolism — from biochemistry to genomics. Nat. Rev. Microbiol. 3:937–947.

Kensche PR, Oti M, Dutilh BE, Huynen MA. 2008. Conservation of divergent transcription in fungi. Trends Genet. 24:207–211.

King N, Rokas A. 2017. Embracing Uncertainty in Reconstructing Early Animal Evolution. Curr. Biol. 27:R1081–R1088.

Kishino H, Miyata T, Hasegawa M. 1990. Maximum likelihood inference of protein phylogeny and the origin of chloroplasts. J. Mol. Evol. 31:151–160.

Kjærbølling I, Vesth TC, Frisvad JC, Nybo JL, Theobald S, Kuo A, Bowyer P, Matsuda Y, Mondo S, Lyhne EK, et al. 2018. Linking secondary metabolites to gene clusters through genome sequencing of six diverse *Aspergillus* species. Proc. Natl. Acad. Sci. 115:E753–E761.

Kobayashi T, Abe K, Asai K, Gomi K, Juvvadi PR, Kato M, Kitamoto K, Takeuchi M, Machida M. 2007. Genomics of *Aspergillus oryzae*. Biosci. Biotechnol. Biochem. 71:646–670.

Kobert K, Salichos L, Rokas A, Stamatakis A. 2016. Computing the Internode Certainty and Related Measures from Partial Gene Trees. Mol. Biol. Evol. 33:1606–1617.

Kocsubé S, Perrone G, Magistà D, Houbraken J, Varga J, Szigeti G, Hubka V, Hong S-B, Frisvad JC, Samson RA. 2016. *Aspergillus* is monophyletic: Evidence from multiple gene phylogenies and extrolites profiles. Stud. Mycol. 85:199–213.

Kolde R. 2012. Package ‘pheatmap’. Bioconductor:1–6.

Kosiol C, Goldman N. 2005. Different Versions of the Dayhoff Rate Matrix. Mol. Biol. Evol. 22:193–199.

Kumar S, Filipski AJ, Battistuzzi FU, Kosakovsky Pond SL, Tamura K. 2012. Statistics and Truth in Phylogenomics. Mol. Biol. Evol. 29:457–472.

Da Lage J-L, Binder M, Hua-Van A, Janeček Š, Casane D. 2013. Gene make-up: rapid and massive intron gains after horizontal transfer of a bacterial α-amylase gene to Basidiomycetes. BMC Evol. Biol. 13:40.

Lê S, Josse J, Husson F. 2008. FactoMineR?: An R Package for Multivariate Analysis. J. Stat. Softw. 25:1–18.

Le SQ, Gascuel O. 2008. An Improved General Amino Acid Replacement Matrix. Mol. Biol. Evol. 25:1307–1320.

Li B, Zong Y, Du Z, Chen Y, Zhang Z, Qin G, Zhao W, Tian S. 2015. Genomic Characterization Reveals Insights Into Patulin Biosynthesis and Pathogenicity in *Penicillium* Species. Mol. Plant-Microbe Interact. 28:635–647.

Li Y, Liu Z, Shi P, Zhang J. 2010. The hearing gene Prestin unites echolocating bats and whales. Curr. Biol. 20:R55–R56.

Liu L, Zhang J, Rheindt FE, Lei F, Qu Y, Wang Y, Zhang Y, Sullivan C, Nie W, Wang J, et al. 2017. Genomic evidence reveals a radiation of placental mammals uninterrupted by the KPg boundary. Proc. Natl. Acad. Sci. 114:E7282–E7290.

Ma L-J, Geiser DM, Proctor RH, Rooney AP, O’Donnell K, Trail F, Gardiner DM, Manners JM, Kazan K. 2013. *Fusarium* Pathogenomics. Annu. Rev. Microbiol. 67:399–416.

Macheleidt J, Mattern DJ, Fischer J, Netzker T, Weber J, Schroeckh V, Valiante V, Brakhage AA. 2016. Regulation and Role of Fungal Secondary Metabolites. Annu. Rev. Genet. 50:371–392.

Madden T. 2013. The BLAST sequence analysis tool. BLAST Seq. Anal. Tool:1–17.

Magan N, Lacey J. 1984. Effects of gas composition and water activity on growth of field and storage fungi and their interactions. Trans. Br. Mycol. Soc. 82:305–314.

Marcet-Houben M, Ballester A-R, de la Fuente B, Harries E, Marcos JF, González-Candelas L, Gabaldón T. 2012. Genome sequence of the necrotrophic fungus *Penicillium digitatum*, the main postharvest pathogen of citrus. BMC Genomics 13:646.

Marín, Sanchis, Sáenz, Ramos, Vinas, Magan. 1998. Ecological determinants for germination and growth of some *Aspergillus* and *Penicillium* spp. *from maize grain*. J. Appl. Microbiol. 84:25–36.

Martin SH, Dasmahapatra KK, Nadeau NJ, Salazar C, Walters JR, Simpson F, Blaxter M, Manica A, Mallet J, Jiggins CD. 2013. Genome-wide evidence for speciation with gene flow in *Heliconius* butterflies. Genome Res. 23:1817–1828.

Mead ME, Knowles SL, Raja HA, Beattie SR, Kowalski CH, Steenwyk JL, Silva LP, Chiaratto J, Ries LNA, Goldman GH, et al. 2018. Characterizing the pathogenic, genomic, and chemical traits of *Aspergillus fischeri*, the closest sequenced relative of the major human fungal pathogen *Aspergillus fumigatus*. bioRxiv.

Mirarab S, Warnow T. 2015. ASTRAL-II: coalescent-based species tree estimation with many hundreds of taxa and thousands of genes. Bioinformatics 31:i44–i52.

Morgulis A, Coulouris G, Raytselis Y, Madden TL, Agarwala R, Schäffer AA. 2008. Database indexing for production MegaBLAST searches. Bioinformatics 24:1757–1764.

Mount DW. 2008. Using BLOSUM in Sequence Alignments. Cold Spring Harb. Protoc. 2008:pdb.top39-pdb.top39.

Nabhan AR, Sarkar IN. 2012. The impact of taxon sampling on phylogenetic inference: a review of two decades of controversy. Brief. Bioinform. 13:122–134.

Nguyen L-T, Schmidt HA, von Haeseler A, Minh BQ. 2015. IQ-TREE: A Fast and Effective Stochastic Algorithm for Estimating Maximum-Likelihood Phylogenies. Mol. Biol. Evol. 32:268–274.

Nielsen JC, Grijseels S, Prigent S, Ji B, Dainat J, Nielsen KF, Frisvad JC, Workman M, Nielsen J. 2017. Global analysis of biosynthetic gene clusters reveals vast potential of secondary metabolite production in *Penicillium* species. Nat. Microbiol. 2:17044.

Nierman WC, Pain A, Anderson MJ, Wortman JR, Kim HS, Arroyo J, Berriman M, Abe K, Archer DB, Bermejo C, et al. 2005. Genomic sequence of the pathogenic and allergenic filamentous fungus *Aspergillus fumigatus*. Nature 438:1151–1156.

O’Donnell K, Rooney AP, Proctor RH, Brown DW, McCormick SP, Ward TJ, Frandsen RJN, Lysøe E, Rehner SA, Aoki T, et al. 2013. Phylogenetic analyses of RPB1 and RPB2 support a middle Cretaceous origin for a clade comprising all agriculturally and medically important fusaria. Fungal Genet. Biol. 52:20–31.

Ogundero VW. 1983. Factors affecting growth and cellulose hydrolysis by the thermotolerant *Aspergillus nidulans* from composts. Acta Biotechnol. 3:65–72.

Patterson N, Richter DJ, Gnerre S, Lander ES, Reich D. 2006. Genetic evidence for complex speciation of humans and chimpanzees. Nature 441:1103–1108.

Phillips MJ, Delsuc F, Penny D. 2004. Genome-Scale Phylogeny and the Detection of Systematic Biases. Mol. Biol. Evol. 21:1455–1458.

Phillips MJ, Penny D. 2003. The root of the mammalian tree inferred from whole mitochondrial genomes. Mol. Phylogenet. Evol. 28:171–185.

Pitt JI. 1994. The current role of *Aspergillus* and *Penicillium* in human and animal health. Med. Mycol. 32:17–32.

Pitt JI. 2002. Biology and Ecology of Toxigenic *Penicillium* Species. In: Advances in experimental medicine and biology. Vol. 504. p. 29–41.

Pitt JI, Hocking AD. 2009. Fungi and Food Spoilage. Boston: Springer

Pitt JI, Taylor JW. 2014. *Aspergillus*, its sexual states and the new International Code of Nomenclature. Mycologia 106:1051–1062.

R Development Core Team R. 2008. Computational Many-Particle Physics. (Fehske H, Schneider R, Weiße A, editors.). Berlin, Heidelberg: Springer Berlin Heidelberg

Raftery AE, Lewis SM. 1995. The number of iterations, convergence diagnostics and generic Metropolis algorithms. Pract. Markov Chain Monte Carlo 7:763–773.

Rambaut A. 2009. FigTree, a graphical viewer of phylogenetic trees. Inst. Evol. Biol. Univ. Edinburgh.

Dos Reis M, Yang Z. 2013. The unbearable uncertainty of Bayesian divergence time estimation. J. Syst. Evol. 51:30–43.

Rohlfs M, Albert M, Keller NP, Kempken F. 2007. Secondary chemicals protect mould from fungivory. Biol. Lett. 3:523–525.

Rohlfs M, Churchill ACL. 2011. Fungal secondary metabolites as modulators of interactions with insects and other arthropods. Fungal Genet. Biol. 48:23–34.

Rokas A, Carroll SB. 2005. More Genes or More Taxa? The Relative Contribution of Gene Number and Taxon Number to Phylogenetic Accuracy. Mol. Biol. Evol. 22:1337–1344.

Rokas A, Williams BL, King N, Carroll SB. 2003. Genome-scale approaches to resolving incongruence in molecular phylogenies. Nature 425:798–804.

Rokas A, Wisecaver JH, Lind AL. 2018. The birth, evolution and death of metabolic gene clusters in fungi. Nat. Rev. Microbiol.

Ropars J, Cruaud C, Lacoste S, Dupont J. 2012. A taxonomic and ecological overview of cheese fungi. Int. J. Food Microbiol. 155:199–210.

Salichos L, Rokas A. 2013. Inferring ancient divergences requires genes with strong phylogenetic signals. Nature 497:327–331.

Salichos L, Stamatakis A, Rokas A. 2014. Novel Information Theory-Based Measures for Quantifying Incongruence among Phylogenetic Trees. Mol. Biol. Evol. 31:1261–1271.

Samson RA, Visagie CM, Houbraken J, Hong S-B, Hubka V, Klaassen CHW, Perrone G, Seifert KA, Susca A, Tanney JB, et al. 2014. Phylogeny, identification and nomenclature of the genus *Aspergillus*. Stud. Mycol. 78:141–173.

Sang T, Zhong Y. 2000. Testing Hybridization Hypotheses Based on Incongruent Gene Trees.Olmstead R, editor. Syst. Biol. 49:422–434.

Sayers EW, Barrett T, Benson DA, Bryant SH, Canese K, Chetvernin V, Church DM, DiCuccio M, Edgar R, Federhen S, et al. 2009. Database resources of the National Center for Biotechnology Information. Nucleic Acids Res. 37:D5–D15.

Sedgwick P. 2014. Spearman’s rank correlation coefficient. BMJ 349:g7327–g7327.

Sharpton TJ, Stajich JE, Rounsley SD, Gardner MJ, Wortman JR, Jordar VS, Maiti R, Kodira CD, Neafsey DE, Zeng Q, et al. 2009. Comparative genomic analyses of the human fungal pathogens *Coccidioides* and their relatives. Genome Res. 19:1722–1731.

Shen X-X, Hittinger CT, Rokas A. 2017. Contentious relationships in phylogenomic studies can be driven by a handful of genes. Nat. Ecol. Evol. 1:0126.

Shen X-X, Salichos L, Rokas A. 2016. A Genome-Scale Investigation of How Sequence, Function, and Tree-Based Gene Properties Influence Phylogenetic Inference. Genome Biol. Evol. 8:2565–2580.

Shen X-X, Zhou X, Kominek J, Kurtzman CP, Hittinger CT, Rokas A. 2016. Reconstructing the Backbone of the Saccharomycotina Yeast Phylogeny Using Genome-Scale Data. G3 Genes|Genomes|Genetics 6:3927–3939.

Shimodaira H. 2002. An Approximately Unbiased Test of Phylogenetic Tree Selection. Syst. Biol. 51:492–508.

Shimodaira H, Hasegawa M. 1999. Multiple Comparisons of Log-Likelihoods with Applications to Phylogenetic Inference. Mol. Biol. Evol. 16:1114.

Song S, Liu L, Edwards S V., Wu S. 2012. Resolving conflict in eutherian mammal phylogeny using phylogenomics and the multispecies coalescent model. Proc. Natl. Acad. Sci. 109:14942–14947.

Squire R. 1981. Ranking animal carcinogens: a proposed regulatory approach. Science (80-. ). 214:877–880.

Stamatakis A. 2014. RAxML version 8: a tool for phylogenetic analysis and post-analysis of large phylogenies. Bioinformatics 30:1312–1313.

Stanke M, Waack S. 2003. Gene prediction with a hidden Markov model and a new intron submodel. Bioinformatics 19:ii215–ii225.

Stierle AA, Stierle DB. 2015. Bioactive Secondary Metabolites Produced by the Fungal Endophytes of Conifers. Nat. Prod. Commun. 10:1671–1682.

Suh A. 2016. The phylogenomic forest of bird trees contains a hard polytomy at the root of Neoaves. Zool. Scr. 45:50–62.

Tavaré S. 1986. Some probabilistic and statistical problems in the analysis of DNA sequences. Lect. Math. life Sci. 17:57–86.

Taylor JW, Göker M, Pitt JI. 2016. Choosing one name for pleomorphic fungi: The example of *Aspergillus* versus *Eurotium*, *Neosartorya* and *Emericella*. Taxon 65:593–601.

Vijaykrishna D, Jeewon R, Hyde K. 2006. Molecular taxonomy, origins and evolution of freshwater ascomycetes. Fungal Divers. 23:351–390.

Vinet L, Zhedanov A. 2011. A ‘missing’ family of classical orthogonal polynomials. J. Phys. A Math. Theor. 44:085201.

Vinnere Pettersson O, Leong SL. 2011. Fungal Xerophiles (Osmophiles). In: eLS. Chichester, UK: John Wiley & Sons, Ltd.

de Vries RP, Riley R, Wiebenga A, Aguilar-Osorio G, Amillis S, Uchima CA, Anderluh G, Asadollahi M, Askin M, Barry K, et al. 2017. Comparative genomics reveals high biological diversity and specific adaptations in the industrially and medically important fungal genus *Aspergillus*. Genome Biol. 18:28.

Waddell PJ, Steel M. 1997. General Time-Reversible Distances with Unequal Rates across Sites: Mixing Γ and Inverse Gaussian Distributions with Invariant Sites. Mol. Phylogenet. Evol. 8:398–414.

Waterhouse RM, Seppey M, Simão FA, Manni M, Ioannidis P, Klioutchnikov G, Kriventseva E V, Zdobnov EM. 2018. BUSCO Applications from Quality Assessments to Gene Prediction and Phylogenomics. Mol. Biol. Evol. 35:543–548.

Waterhouse RM, Tegenfeldt F, Li J, Zdobnov EM, Kriventseva E V. 2013. OrthoDB: a hierarchical catalog of animal, fungal and bacterial orthologs. Nucleic Acids Res. 41:D358–D365.

Wickham H. 2009. ggplot2. New York, NY: Springer New York

Yang Y, Chen M, Li Z, Al-Hatmi AMS, de Hoog S, Pan W, Ye Q, Bo X, Li Z, Wang S, et al. 2016. Genome Sequencing and Comparative Genomics Analysis Revealed Pathogenic Potential in *Penicillium capsulatum* as a Novel Fungal Pathogen Belonging to Eurotiales. Front. Microbiol. 7.

Yang Z. 1994. Maximum likelihood phylogenetic estimation from DNA sequences with variable rates over sites: Approximate methods. J. Mol. Evol. 39:306–314.

Yang Z. 1996. Among-site rate variation and its impact on phylogenetic analyses. Trends Ecol. Evol. 11:367–372.

Yang Z. 2007. PAML 4: Phylogenetic Analysis by Maximum Likelihood. Mol. Biol. Evol. 24:1586–1591.

Zhong B, Liu L, Yan Z, Penny D. 2013. Origin of land plants using the multispecies coalescent model. Trends Plant Sci. 18:492–495.

Zhou X, Lutteropp S, Czech L, Stamatakis A, Looz M von, Rokas A. 2017. Quartet-based computations of internode certainty provide accurate and robust measures of phylogenetic incongruence. bioRxiv:168526.

Zhou X, Peris D, Kominek J, Kurtzman CP, Hittinger CT, Rokas A. 2016. in silico Whole Genome Sequencer & Analyzer (iWGS): A Computational Pipeline to Guide the Design and Analysis of de novo Genome Sequencing Studies. G3 Genes|Genomes|Genetics.

Zhou X, Shen X-X, Hittinger CT, Rokas A. 2018. Evaluating Fast Maximum Likelihood-Based Phylogenetic Programs Using Empirical Phylogenomic Data Sets. Mol. Biol. Evol. 35:486–503.

